# Genome-wide association study reveals complex genetic architecture of cadmium and mercury accumulation and tolerance traits in *Medicago truncatula*

**DOI:** 10.1101/2021.10.29.465169

**Authors:** Tim Paape, Benjamin Heiniger, Miguel Santo Domingo, Michael R. Clear, M. Mercedes Lucas, José J. Pueyo

## Abstract

Heavy metals are an increasing problem due to contamination from human sources that and can enter the food chain by being taken up by plants. Understanding the genetic basis of accumulation and tolerance in plants is important for reducing the uptake of toxic metals in crops and crop relatives, as well as for removing heavy metals from soils by means of phytoremediation. Following exposure of *Medicago truncatula* seedlings to cadmium (Cd) and mercury (Hg), we conducted a genome-wide association study using relative root growth (RRG) and leaf accumulation measurements. Cd and Hg accumulation and RRG had heritability ranging 0.44 - 0.72 indicating high genetic diversity for these traits. The Cd and Hg trait associations were broadly distributed throughout the genome, indicated the traits are polygenic and involve several quantitative loci. For all traits, candidate genes included several membrane associated ATP-binding cassette transporters, P-type ATPase transporters, several oxidative stress response genes and stress related UDP-glycosyltransferases. The P-type ATPase transporters and ATP-binding cassette protein-families have roles in vacuole transport of heavy metals, and our findings support their wide use in physiological plant responses to heavy metals and abiotic stresses. We also found associations between Cd RRG with the genes *CAX3* and *PDR3*, two linked adjacent genes, and leaf accumulation of Hg associated with the genes *NRAMP6* and *CAX9*. When plant genotypes with the most extreme phenotypes were compared, we found significant divergence in genomic regions using population genomics methods that contained metal transport and stress response gene ontologies. Several of these genomic regions show high linkage disequilibrium (LD) among candidate genes suggesting they have evolved together. Minor allele frequency (MAF) and effect size of the most significant SNPs was negatively correlated with large effect alleles being most rare. This is consistent with purifying selection against alleles that increase toxicity and abiotic stress. Conversely, the alleles with large affect that had higher frequencies that were associated with the exclusion of Cd and Hg. Overall, macroevolutionary conservation of heavy metal and stress response genes is important for improvement of forage crops by harnessing wild genetic variants in gene banks such as the Medicago HapMap collection.

## Introduction

Heavy metals are high-density elements that can cause toxic effects when present in excess quantities. Cadmium (Cd) and mercury (Hg) are two of the most toxic heavy metals to humans as Cd poisoning can cause kidney damage and osteoporosis (Järup and Åkesson, 2009), while Hg poisoning is associated with lung, kidney, muscle and brain damage (Vallee and Ulmer, 1972; Bernhoft, 2012). Heavy metals may occur naturally at low concentrations in soils, often originating from volcanic soils and weathered rocks. However, the predominant source of heavy metal contamination are mines, foundries, and smelters, which are often associated with particularly high levels of contamination in surrounding soils (Tchounwou et al., 2012; Alloway, 2013). This pollution can negatively affect surrounding agricultural and natural ecosystems. Heavy metal accumulation in agricultural soils is worsened by atmospheric deposition, sewage irrigation practices, and the extensive use of soil amendments, livestock manures, pesticides and agrophytochemicals (Nicholson et al., 2003; Peng et al., 2019). Plants grown on contaminated soils may accumulate heavy metals in aerial parts such as leaf tissues and seeds and can result in severe health consequences for foraging animals and humans if these metals enter the food supply (Peralta-Videa et al., 2009).

In plants, heavy metals such as Cd and Hg are non-essential ions that can be taken up from the environment by essential micronutrient transporters (Clemens, 2006) and subsequently assimilated into the aerial parts of the plants via the roots. High intracellular accumulation of metal ions can lead to the denaturation of proteins, the displacement of essential metals from biomolecules, problems in membrane integrity, and the formation of reactive oxygen species (ROS). Excess metals in leaves can result in chlorosis, disruption of photosynthetic pathways, and breakdown of basic metabolic processes often leading to the death of the plant. Managing heavy metal toxicity by plants requires several genetic loci for integrated transport and tissue detoxification that operate at both the cellular and molecular level. Thus, traits associated with tolerance and accumulation are expected to be polygenic.

Molecular studies have shown that plants bind Cd and Hg with phytochelatins that are synthesized from glutathione and cysteine through the activation of the phytochelatin synthase (PCS) enzymatic pathway (Cobbett, 2001; Hossain et al., 2012). Binding and chelation of Cd and Hg ions enable their transport across membranes where they are sequestered into vacuoles for sub-cellular compartmentalization (Martinoia et al., 2018). The ATP-binding cassette subfamily of transporters (ABC-transporters), specifically *ABCC1* and *ABCC2* in *A. thaliana* and poplar, transport chelated Cd and Hg ions into vacuoles (Park et al., 2012; Brunetti et al., 2015; Sun et al., 2018) which reduces cellular toxicity and limits dissemination throughout the plant. Additionally, unchelated Cd ions can be directly sequestered into the vacuole and mediated by the heavy metal ATPase *HMA3* (Morel et al., 2008; Chao et al., 2012) and the cation exchange (CAX) type antiporters such as *AtCAX2* and *AtCAX4* (Cheng et al., 2005; Punshon et al., 2012). In rice (*Oryza sativa*), the natural resistance-associated macrophage protein 5 (*NRAMP5*, also a manganese (Mn) transporter), is the main transporter of Cd into roots through the apoplast (Sasaki et al., 2012; Clemens and Ma, 2016). *NRAMP6* also plays a role in Cd accumulation in *A. thaliana* as mutants confer greater Cd tolerance (Cailliatte et al., 2009). Translocation of heavy metals from the root into the shoot occurs by loading ions into the xylem often using heavy metal ATPases (HMA’s), such as *HMA2* and *HMA4*, in *A. thaliana* and rice (Hanikenne et al., 2008; Takahashi et al., 2012). While far less is known about intracellular transport of Hg, there may be conservation between Cd and Hg transporters at the protein-family level given the similar vacuole transport mechanism for both Cd and Hg in *A. thaliana* using the same ABC-transporters (Park et al., 2012).

Identifying the physiological processes and genetic mechanisms that plants use to limit the accumulation of toxic heavy metals is important for human health and agriculture. The wild relatives of crop species possess standing variation that can be useful for identifying the molecular mechanisms associated with heavy metal tolerance and accumulation (Chao et al., 2012; Wu et al., 2015; Zhao et al., 2017; Chen et al., 2018). For example, allelic variants for reduced accumulation of heavy metals (excluder alleles) can minimize toxicity in plants grown in natural and agricultural ecosystems. Mining these genetic variants in species-wide germplasm collections of crop relatives can help us identify single nucleotide polymorphisms (SNPs) for both excluder and accumulator alleles and their associated genes. Conversely, accumulator alleles in highly tolerant plant genotypes may possess useful genetic adaptations that can facilitate phytoremediation of toxic soils (Swartjes, 2011; Yan et al., 2020) by increasing the capacity of plants to absorb high levels of toxic metals.

We are interested in the genetic architecture of tolerance to Cd and Hg and leaf accumulation of these metals in the model legume species *Medicago truncatula*, which is a Mediterranean forage legume and close relative of important crop legumes. Legumes such as *M. truncatula* have evolved symbiotic interactions with nitrogen-fixing bacteria (rhizobia) that can enrich plants with nitrogen. Symbiosis begins when the roots of the host plant come into contact with soil-borne rhizobia, resulting in nodule formation on the roots (Kevei et al., 2002; Van de Velde et al., 2010). Inside the nodules, the rhizobia fix nitrogen for the host-plant in exchange for carbon nutrients from host-plants to rhizobia. Therefore, the ability of the host-plant to detoxify plant tissues, particularly roots, is essential for rhizobia to infect roots and form nodules (León-Mediavilla et al., 2018). Detoxification of roots involves both compartmentalization of toxic ions (i.e., in vacuoles) and transport of the ions away from roots often into aerial tissues such as leaves. However, excessive transport of toxic ions to aerial tissues would come at a high cost to plant fitness due to physiological stress and disruption of photosynthesis.

Considerable genomic resources exist for *M. truncatula* including high quality reference genomes and gene annotations (Young et al., 2011; Tang et al., 2014), genome-wide polymorphism data (Branca et al., 2011; Paape et al., 2013), a HapMap panel of resequenced germplasm for conducting genome wide association studies (GWAS) (Stanton-Geddes et al., 2013; Bonhomme et al., 2014; Kang et al., 2019), and a large mutant collection (Lee et al., 2018). These resources allow us to map phenotypes to very fine genomic regions and identify candidate genes associated with traits of interest, providing the basis for functional genetics studies (Curtin et al., 2017). Moreover, in previous studies Hg- and Cd-sensitive and tolerant cultivars were identified by phenotyping *M. truncatula* germplasm resources, and differential responses to metal stress between tolerant and sensitive cultivars were reported (García de la Torre et al., 2013, 2021). Both cultivar selection and transgenic approaches can be used to obtain legumes with increased tolerance to abiotic stress (Coba de la Peña and Pueyo, 2012). Moreover, tolerant rhizobia isolated from contaminated soils (Nonnoi et al., 2012) or genetically modified (Shvaleva et al., 2010) can be used as inoculants to increase legume tolerance to heavy metal stress (Quiñones et al., 2013; Arregui et al., 2021).

Standing genetic variation for quantitative traits, such as phenotypic responses to heavy metals, is useful for detecting allelic variation (i.e., SNPs) or genomic divergence associated with trait variation. The variation present in species-wide samples from natural populations (such as the Medicago HapMap panel) may be comprised of adaptive and deleterious alleles that have been tested in highly variable, natural habitats. In addition to identifying genetic loci associated with traits, GWAS can be used to estimate allele frequencies of associated SNPs and their effect size, which may reveal the forces of selection that contributed to the genetic architecture of a trait (Stinchcombe and Hoekstra, 2007; Josephs et al., 2017). In GWAS data, we expect that mutations (SNPs) with larger effect sizes will most often be deleterious (Eyre-Walker and Keightley, 2007) and be subject to purifying or negative selection (Trotter, 2014) under mutation-selection balance (Crow and Kimura, 2010). These processes can be revealed by negative correlations between SNP effect size (estimated in the GWAS) and minor allele frequencies (MAFs), where SNPs with smaller effect on a trait will be at higher frequency and those with a larger effect will be at lower frequencies (Stanton-Geddes et al., 2013; Josephs et al., 2015). In addition, the phenotypic distributions exhibited by standing variation may also correspond to genomic divergence. By treating genotypic groups at opposite ends of phenotypic distributions as populations, it may be possible to detect signals of genomic divergence using population differentiation methods such as F-statistics (F_st_) and composite likelihood ratio (CLR) tests. F-statistics are traditionally used to test for natural selection between populations by comparing allele frequencies within and between groups (Holsinger and Weir, 2009), while CLR tests detect selective sweeps between two populations (XP-CLR; Chen et al., 2010). These tests may provide complementary information to GWAS regarding genomic regions underlying divergent trait values.

Toxic levels of Cd and Hg in plants can be estimated based on relative root growth (RRG) and leaf accumulation. RRG acts as a measure of tolerance or root growth inhibition (García de la Torre et al., 2013, 2021) while leaf accumulation measured using ionomics characterizes leaf-level responses such as root-to-shoot transport and leaf capacity. In this study we set out to (1) quantify the phenotypic variation of Cd and Hg tolerance and accumulation in the Medicago HapMap panel, (2) identify genes that are associated with these traits using complementary methods to locate genomic regions, (3) characterize the genetic architecture of tolerance and accumulation of these two metals, and (4) estimate whether the genetic architecture has been shaped by selection acting upon natural variation. When orthologous genes are detected in GWAS experiments across multiple plant species it suggests conservation of metal transporters and detoxification mechanisms across the plant kingdom. Overall, the expected results of the present study will contribute to our understanding of the mechanisms involved in heavy metal tolerance, translocation and accumulation in plants, and will provide candidate targets to modify heavy metal accumulation or tolerance by traditional breeding or transgenic approaches.

## Methods

### Plant Growth and Measurements of Phenotypes

Seeds from 236 resequenced *M. truncatula* genotypes were obtained from the University of Minnesota, Medicago HapMap project (http://www.medicagohapmap.org/hapmap/germplasm). Two separate heavy metal treatments were applied in parallel to the set of *M. truncatula* HapMap genotypes at the seedling stage. A modified Hoagland nutrient solution (based on García de la Torre et al., 2013; 2021) was used to grow plants in hydroponic conditions using the following nutrient concentrations: 2.02 gL^-1^ KNO_3_, 0.68gL^-1^ KH_2_PO_4_, 0.182gL^-1^ CaCl_2_·2H_2_O, 0.615gL^-1^ MgSO_4_·7H_2_O, 0.109 gL^-1^ K_2_SO_4_, 0.205 gL^-1^ Hampiron (Rhône Poulenc), and 1.35 mL of a solution containing: 11 gL^-1^ H_3_BO_3_, 6.2 gL^-1^ MnSO_4_ ·H_2_O, 10 gL^-1^ KCl, 1 gL^-1^ ZnSO_4_ ·7H_2_O, 1 gL^-1^ (NH_4_)_6_ Mo_7_ O_24_ ·4H_2_O, 0.5 gL^-1^ CuSO_4_·5H_2_O and 0.5mLL^-1^ H_2_SO_4_. One set of plants was treated with mercury using 4μM HgCl_2_ added to the Hoagland solution. A second set of plants was treated with cadmium using 10μM CdCl_2_ added to the Hoagland solution. A third set of plants was given no heavy metal treatment and was used as a control group. Each treatment began with fifteen replicate plant seedlings for each *M. truncatula* genotype. Four traits were measured on the *M. truncatula* Hapmap panel following the heavy metal treatments: 1) Relative Root Growth (RRG) in plants treated with Hg and 2) RRG in plants treated with Cd, 3) accumulation of Cd in leaf tissues, 4) and accumulation of Hg in leaf tissues. Seedlings were placed for 24 h in the hydroponic system with 250 mL of nutrient solution for acclimatizing in growth chamber conditions (24/20ºC, 16/8h photoperiod), and then 48 h in the same conditions with or without the heavy metal treatment. RRG is a reliable indicator of metal tolerance in *M. truncatula* (García de la Torre et al., 2013, 2021). Root length was measured at 24 h of plants growing in the hydroponic medium before metal was added, and 48 h later, after applying the metal treatments. Photo images of the seedlings were taken at 24 h and again after the 48 h treatment. Roots were measured using the software ImageJ. For calculating the RRG of the seedlings, the increase in length is normalized by the increase in control seedlings (Equation 1):

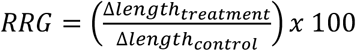

For metal concentration measurements in leaves, cotyledons were harvested after plants had been exposed to Cd or Hg treatment for 48h. The leaf tissues were washed with 10 mM Na_2_EDTA to remove traces of metals on their surface. Washed and dried leaves were digested using concentrated HNO_3_ and H_2_O_2_. After the tissue digestion, distilled water was added, then the mixture was filtered. Cd and Hg concentrations were measured using inductively coupled plasma atomic emission spectroscopy (ICP-AES). For leaf accumulation, three samples (biological replicates) were analyzed per genotype. We used R version 3.3.3 to conduct statistical on phenotypic distributions and the lmer4 package to estimate variance components to calculate broad sense heritability (*H*^2^). Metal concentrations were determined as previously reported (García de la Torre et al., 2013, 2021).

### Genome-wide Association Study

Genome-wide SNP data (BCF files) called against the Mt4.0 version of the reference genome from 262 *Medicago truncatula* accessions was obtained from the Medicago HapMap project (http://www.medicagohapmap.org/downloads/mt40). Missing SNPs (due to low coverage in some genotypes) were imputed using BEAGLE version 4.1 (Browning and Browning, 2016) with default parameters. The imputed dataset was split by chromosome and converted to hapmap format (.hmp) using TASSEL 5 (Bradbury et al., 2007). We used the software Admixture (Alexander et al., 2009) to generate a population structure co-variance matrix. The following criteria was used to select independent SNPs for the Admixture analysis, based on similar criteria used by Gentzbittel *et al*. (2019) to remove SNPs that were linked or below allele frequency thresholds. SNPs were filtered by genotyping rate and minor allele frequency, then converted to bed format using PLINK 1.9 beta 6.5 (Chang et al., 2015) using the command line input parameters: --geno 0.05 −maf 0.01 --make-bed. Independent sites were extracted using PLINK using the following input parameters: -indep 300 60 1.22. Admixture v1.3.0 was run on 843,307 SNPs and k values ranging from 1 to 10 were estimated by running 10 iterations per K with a different seed value for each iteration (--seed=1 to --seed=10). For each k, average cross validation errors were calculated and the iteration with lowest CV error was plotted in R. The SNP dataset containing 262 HapMap accessions was filtered to contain accessions with phenotype data present in the current study, resulting in a SNP dataset containing 236 genotypes.

We ran the GWAS using GAPIT (Lipka et al., 2012; Tang et al., 2016) version 20160323, once with the population structure covariance matrix (with lowest cross validation error, k = 5), and a second time without the structure covariance matrix. A kinship (K) matrix, to control for genetic relatedness was estimated by the software. The parameters used in the GAPIT model were KI = NULL, PCA.total = 3, SNP.MAF = 0.02, SNP.fraction = 0.6, Major.allele.zero = TRUE and Geno.View.output = FALSE. We also ran the GWAS using GEMMA (Zhou and Stephens, 2012). The imputed dataset was filtered to only contain phenotyped individuals with bcftools 1.2 and subsequently converted to bed format using PLINK 2.0 alpha while the phenotype data was converted to fam format using the PLINK option: --make-just-fam. We then used GEMMA 0.98.1 to calculate relatedness matrices for each trait and to perform GWAS iterations using a multivariate linear mixed model and a minor allele frequency cutoff of 2 percent (parameters: -lmm -maf 0.02). As with GAPIT, GEMMA was run with and without a population structure covariance matrix for comparison.

We avoided using strict p-value thresholds because they would be less informative for characterizing the genetic architecture of polygenic traits. Instead, we explored the larger landscape of SNPs by utilizing a range of p-values. Because we began with several million SNPs, the top 100 to 1000 SNPs with the lowest p-values for each trait are potentially relevant based on functional annotations of genes in the vicinity of these SNPs. The 1000 most significant SNPs across all chromosomes were annotated using a custom Python script for the genes within 1 kb distance from these SNPs using the gene context files available from the HapMap project (http://www.medicagohapmap.org). Among these 1000, we can select any subset such as the top 100 for each trait. Information including distance between SNP and gene, or the nucleotide substitution type (i.e., synonymous, missense), were extracted from the gene context files. Additionally, the blastn tool provided by NCBI (Camacho et al., 2009) was used to perform BLAST of the genes against *Arabidopsis thaliana*. Gene descriptions, gene ontologies (GO-terms) and tissue-specific RNAseq expression levels (from six tissue types in the *M. truncatula* A17 genotype) were further annotated using information from MedicMine (Krishnakumar et al., 2015). Initially, genes with relevance to heavy metal tolerance were categorized as ATPase, metal ion, ion transport, and stress, based on their functional annotation in *M. truncatula*, GO-terms, and the annotation of the closest BLAST ortholog in *A. thaliana*. A statistical test for GO-term enrichment using the genes with the 1000 most significant SNPs was conducted using AgriGO (Du et al., 2010) using false discovery adjusted p-value ≤ 0.05. Regions with a high density of candidate SNPs (“GWAS peaks”) were manually identified using IGV 2.6.3 (Robinson et al., 2011; Thorvaldsdóttir et al., 2013). To calculate pairwise linkage disequilibrium (LD) between the 100 most significant SNPs in the peaks, vcftools was used with the –geno-r2 flag and the resulting values were plotted using the R package LDHeatmap (Shin et al., 2006).

### Quantifying genomic divergence using phenotypic divergence

The phenotypic distributions of each trait (**Figure 1, Supplementary Figure S1**) were used to define two groups, high and low, (per trait) based on trait measurements in the opposite tails of the distributions. For each of the traits measured, the two groups consisted of 30 individuals with the lowest phenotype values and the other consisted of 30 individuals with the highest phenotype values. We used *vcftools* (Danecek et al., 2011) to generate F_st_ statistics with the previously mentioned groups and a window and step size of 100 kb. Similarly, xpclr 1.1 (https://github.com/hardingnj/xpclr/) was used to calculate XP-CLR (Chen et al., 2010) with identical window and step size. Genes in the top 2 percent of windows with highest values of F_st_ or XP-CLR were annotated in the same way as the GWAS SNPs and were further tested for GO-term enrichment. For some genomic regions, overlap between GWAS peaks and the two statistics were determined.

**Figure 1.**
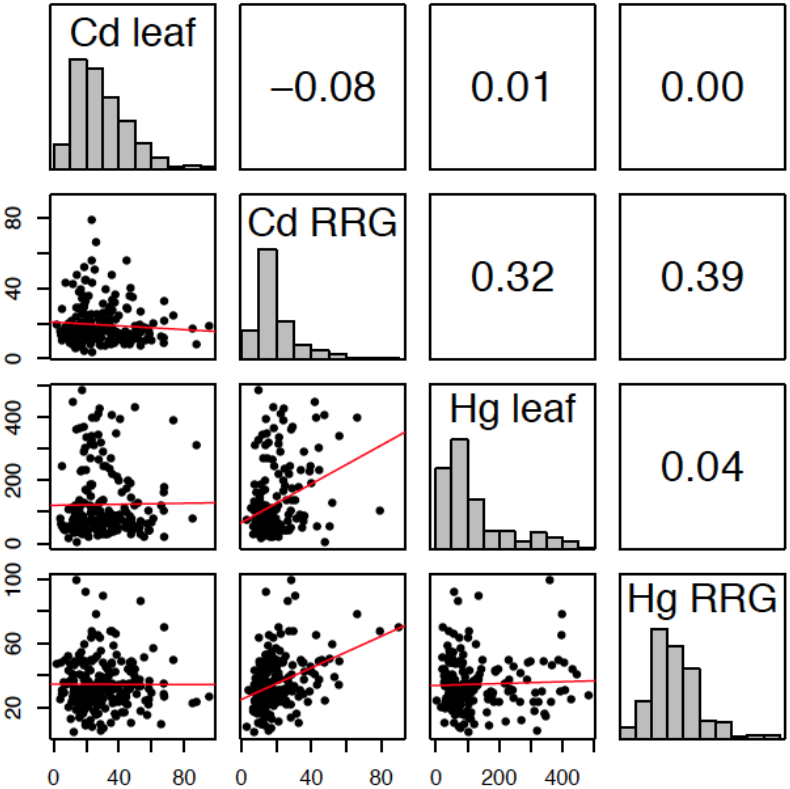
Correlation matrix of Cd leaf accumulation, Cd relative root growth (RRG), Hg leaf accumulation, and Hg relative root growth (RRG) measured in the *M. truncatula* HapMap panel. Distributions of the four phenotypes are shown in the diagonals. The upper, off-diagonal panels contain pairwise Pearson’s correlation coefficients (*r*). Lower, off-diagonals are the plotted trait values of each *M. truncatula* genotype measured in our experiment. The red line shows the slope of the correlation between any pair of traits. The x- and y-axes are values of the phenotype measurements (in μg^-1^ dry weight for leaf, and percent for relative root growth (RRG)).

### Correlation between minor allele frequency and effect size

The correlation between minor allele frequency (MAF) and effect size of the 100 and 1000 most significant GWAS SNPs was determined using Pearson’s correlation coefficient for both GAPIT and GEMMA output datasets. We permuted genotype-phenotype values to make 100 phenotype datasets, then ran 100 iterations of GWAS iterations to make null distributions of MAF and effect sizes. To run the permutations, we used GEMMA, as it performs far better than GAPIT in terms of computational time. For each iteration, Pearson’s correlation coefficients were calculated for minor allele frequency (MAF) and effect size using the 100 and 1000 most significant SNPs. The resulting distribution of correlation coefficients was compared to the correlation of the unpermuted results for each phenotype.

To test whether the most significant SNPs show significant differences than the genomic background, Tajima’s D was calculated for the 1000 most significant SNPs of each trait using *variscan* (Vilella et al., 2005; Hutter et al., 2006) with a window size of 50 bp. We used SNPs that were selected for the Admixture analysis as the background empirical distribution. A t-test was used to determine whether the differences between Tajima’s D of the SNPs identified in the GWAS and those in the genomic background were significant.

## Results

### High Phenotypic Variability, Heritability, and Correlations Among Traits

We measured seedlings from the Medicago HapMap collection for relative root growth (RRG) and metal ion accumulation in leaf tissues following 48 h exposure to Cd and Hg treatments (**Supplementary Table 1**). We found nearly 30-fold difference for Cd RRG and 23-fold difference for Hg RRG between the lowest and highest tolerant genotypes (**Figure 1**). Broad sense heritability (*H*^2^) was equal to 0.61 and 0.72 for the two RRG traits, respectively (**Supplementary Figure 1**). We found 50-fold and 270-fold differences in Cd leaf and Hg leaf accumulation, respectively; *H*^2^ was equal to 0.54 and 0.44 for Cd and Hg leaf accumulation, respectively. No correlation was found between leaf accumulation and RRG for either metal treatment (i.e., plants treated with Cd that had high RRG did not have high leaf accumulation of Cd, and similarly for Hg) (**Figure 1, Supplementary Figure 1**). The highest correlation was between the two root growth traits, Cd and Hg RRG (Pearson’s *r* = 0.39). The heritability estimates and large amount of standing variation suggest that GWAS can identify SNPs and genes underlying these traits due to sufficient genetic differences between genotypes, and that both tolerance and susceptibility alleles are present in the *M. truncatula* HapMap panel.

### Population Structure

We conducted a GWAS to identify genomic regions associated with leaf and root responses to Cd and Hg treatments in *M. truncatula* using a dataset of ∼12 million SNPs after imputing missing data. Among these, we sampled 843k SNPs to generate a covariance matrix (Q) to control for confounding effects of population structure in our GWAS. Based on our analysis of population structure, we determined that k = 5 was the most strongly supported clustering (**Supplementary Figure 2, 3**) using the cross-validation method in Admixture (Alexander et al., 2009). Proportions of each genotype assigned to each of the five clusters were then used in the covariance matrix Q. Including only kinship (K) co-variance in the GWAS had similar empirical and expected distributions of p-values (i.e., similar model fit) in the quantile-quantile plots as did K and Q together (K + Q, kinship plus population structure) for each of the four traits (**Figure 2**). This indicated that kinship largely controlled for confounding genotype relationships with only slight improvement of the model fit by adding structure covariance (e.g., Cd RRG and Hg leaf accumulation). We found that GAPIT and GEMMA performed similarly, but the quantile-quantile plots from GAPIT showed slightly better fitting models (**Figure 2, Supplementary Figure 4**). There were no obvious differences in model fit between either software when the structure covariance matrix was included or excluded. Visual comparison of quantile-quantile plots showed an equal ability of both programs to correct for population structure and kinship.

**Figure 2.**
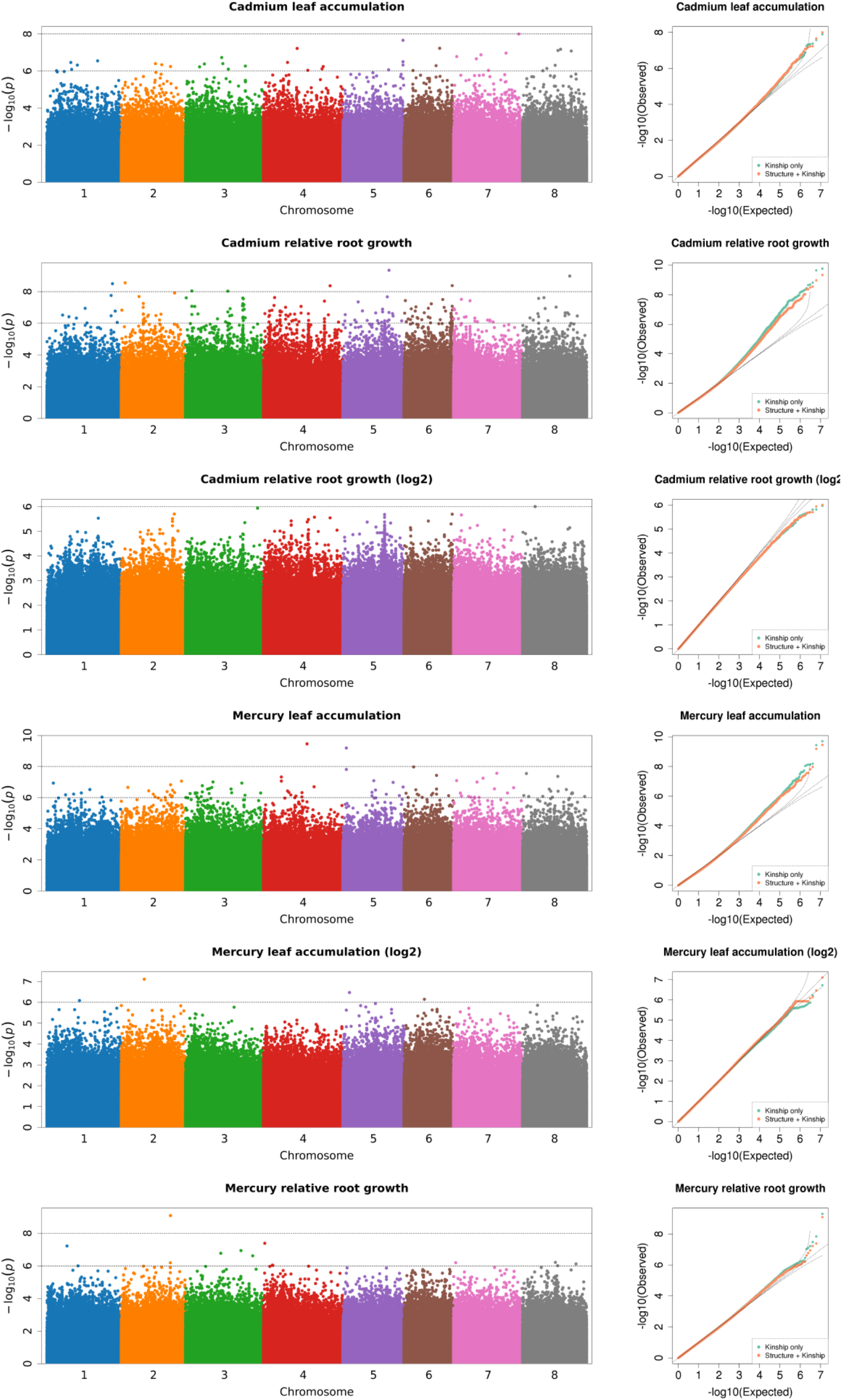
Genome-wide association analysis of heavy metal accumulation and tolerance traits (Manhattan plots shown for each trait; output from GAPIT). Each point represents a SNP at each position in the genome. The number on the x-axis is the chromosome (each color corresponds to one of the eight Mt4.0 chromosomes). The y-axis is the negative log_10_ of the p-values. The higher y-axis values indicate smaller p-values. Next to each Manhattan plot are quantile-quantile (Q-Q) plots which show the model fit with population structure covariance included (orange points), and without population structure covariance (green points). The x-axis of the Q-Q plots is the expected distribution, the y-axis is the empirical distribution.

### Toxic Metal Leaf Accumulation and Relative Root Growth are Polygenic Traits

The genome-wide analyses showed that significant SNPs were widely dispersed across the genome, yet several chromosomal regions associated with each of the four traits were identified (**Figure 2, Figure S4**). This is consistent with our expectation that heavy metal tolerance and metal accumulation are polygenic traits. Candidate gene lists were first generated by selecting the 1000 most significant SNPs (with the lowest p-values, ≤ 1 × 10^−5^; **Supplementary Dataset 1**) within 1 kb proximity of a gene for each trait, which led to about half of all top SNPs being annotated with a gene (SNPs were assumed to be in LD with genes closer than 1 kb, Paape et al., 2012; Paape, 2020). A first pass at identifying candidate genes was done by selecting those with functional annotations based on Blast results to *A. thaliana*, and manual curation of relevant gene annotations and gene ontologies. We first grouped genes with functional annotations for ATPase, ion transport, metal ion, or stress (**Figure 3A**). Among the traits, numbers ranged from 29 - 42 genes that had any of these four functional assignments (**Supplementary Table 2**). The two RRG traits had the most genes, with metal ion and ATPase being the most common in all four of the traits. To determine whether any biological processes or molecular functions were enriched among the candidate genes, we performed tests for gene ontology (GO-term) enrichment for each trait. Response to stress and defense response GO-terms were enriched for Cd RRG, defense response was enriched for Hg leaf, and nucleotide and ATP binding were enriched for Hg RRG (**Supplementary Table 3**). No significant GO-terms were found for Cd leaf.

As a complementary approach to GWAS, we used phenotype values of plant genotypes in the high vs. low ends of the phenotypic distributions, and F_st_ and XP-CLR population genetics statistics to identify genomic regions with high divergence between genotypes within these two groups. We were able to detect significant divergence in genomic regions that contained genes with annotations relevant for heavy metal accumulation and tolerance. The number of genes with ATPase, ion transport, metal ion, or stress related annotations ranged from 69 to 99 total genes per trait (**Figure 3 B, C**). Consistent with the GWAS results, the metal ion category was the most represented across all traits for both population genetics statistics. Among the four traits, GO-term enrichment was most noteworthy for Cd leaf accumulation using the two population genetics methods. For genes identified using F_st_, enriched GO-terms were found for ion binding, cation binding, metal ion binding, response to chemical stimulus, and response to oxidative stress/oxidoreductase activity (**Supplementary Table 4**). For genes identified using XP-CLR, we found GO-term enrichment for oxidoreductase activity, carbon-oxygen lyase activity, FAD binding, and manganese ion binding. The distribution of gene categories and gene ontologies illustrate that by selecting plant genotypes with divergent phenotypes, we were able to detect regions of genomic divergence with biological relevance for metal ion stress.

**Figure 3.**
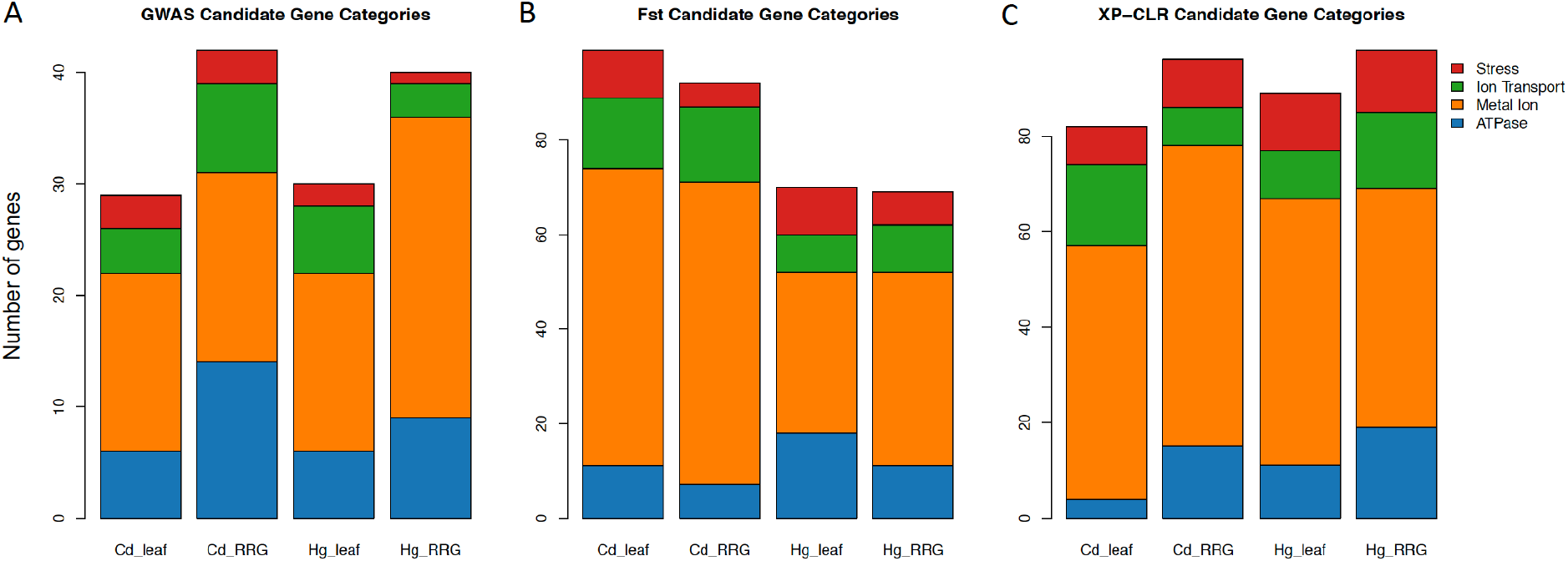
The number of genes with annotations in categories for ATPase, ion transport, metal ion, and stress related genes within 1 kb of the 1000 most significant SNPs using GAPIT for each of the four traits (A). The number of genes in these four categories in the upper 2% of the highest F_st_ sliding windows (B), and the number of genes with these four categories in the 2% of sliding windows with highest XP-CLR values (C). The y-axis represents the number of genes in each category.

### Candidate Gene Identification

To further identify candidate genes detected by GWAS, we looked at gene ontologies among the top 100 SNPs (with the lowest p-values) that mentioned metal ion transport, trans-membrane proteins, and cellular detoxification, and orthologous genes and gene families based on Blast hits to *A. thaliana* that have some role in heavy metal tolerance/accumulation (**Table 1**). For Cd leaf accumulation, among the top 100 SNPs, we found a GTPase membrane protein (Medtr7g009740), a transmembrane amino acid transporter family protein (Medtr8g465310), and an oxidative stress protein, *MtOXS2* (Medtr7g092070). Among the top 1000 significant SNPs for Cd leaf accumulation, was a Fe (II)-dependent oxygenase superfamily protein (Medtr2g069300) with metal ion binding and oxidoreductase activity gene ontologies, and an ATP-binding cassette protein, *MtABCC3* (Medtr5g094830, orthologous to *AtABCC3* (Brunetti et al., 2015)). For Cd RRG, among the top 100 SNPs, we found a non-synonymous SNP in a Fe (II)-dependent oxygenase superfamily protein (Medtr6g090260), a leucine-rich transmembrane protein kinase (Medtr1g100787) with ATPase gene ontology, and a non-synonymous SNP in an ankyrin repeat plant-like protein (Medtr2g438720), which shows highest expression in roots in six tissue-specific comparisons in the A17 ecotype/genotype (**Supplementary Dataset 1**). The top 1000 SNPs for this trait included the ABC-transporters *MtABCC2* (Medtr2g019020) and *MtABCC8* (Medtr8g015980), and Gamma-Glutamyl Transpeptidase 3, *MtGGT3* (Medtr6g090280), which is localized on tonoplasts in *A. thaliana* (Ohkama-Ohtsu et al., 2007), an ortholog of the vacuolar protein *AtVPS45* (Medtr3g090170), and another Fe (II)-dependent oxygenase superfamily protein. We also found associations in the heavy metal transport/detoxification superfamily protein *MtHIPP3* (Medtr2g095480). Heavy-metal-associated Isoprenylated Plant Proteins (HIPP’s) are a family of metallochaperones found only in plants (de Abreu-Neto et al., 2013) that undergo transcriptomic and functional responses to Cd and Zn in *A. thaliana* (Tehseen et al., 2010) and rice (Khan et al., 2019).

**Table 1.** Candidate genes for each trait within 1 kb proximity to top SNPs identified by GWAS. Columns from left to right are chromosome (Chr), Mt4.0 gene ID (Gene Name), SNP position (Position), p-value, proportion of variance explained (PVE) by the SNP, Effect Size (in units of the trait measurement), location or type of nucleotide substitution of a SNP (Effect), *A. thaliana* ortholog identified by BLAST, and gene description. Within each trait, SNPs are sorted smallest to largest p-value.

Cd leaf
| Chr | Gene Name | Position | p-value | PVE | Effect Size | location | BLAST *A. thaliana* | Gene Description |
| --- | --- | --- | --- | --- | --- | --- | --- | --- |
| 7 | Medtr7g009740 | 2221211 | 1.68E-07 | 0.14 | 20.7 | intron | AT5G39900.1 | Small GTP-binding protein |
| 8 | Medtr8g465310 | 23246354 | 4.94E-07 | 0.13 | 20.0 | synonymous | AT3G28960.1 | Transmembrane amino acid transporter family protein |
| 7 | Medtr7g092070 | 36458974 | 1.58E-06 | 0.11 | 19.6 | downstream | AT2G41900.1 | OXS2, OXIDATIVE STRESS 2 |
| 2 | Medtr2g069300 | 28782658 | 5.22E-05 | 0.08 | 10.2 | 3 prime UTR | AT1G06620.1 | Fe(II)-dependent oxygenase superfamily protein |
| 5 | Medtr5g094830 | 41452129 | 3.34E-05 | 0.08 | 16.8 | synonymous | AT3G13080.1 | ABCC3; transmembrane ATPase activity |

Cd RRG
| Chr | Gene Name | Position | p-value | PVE | Effect Size | location | BLAST *A. thaliana* | Gene Description |
| --- | --- | --- | --- | --- | --- | --- | --- | --- |
| 6 | Medtr6g090260 | 34282023 | 4.18E-09 | 0.16 | 17.2 | non-synonymous | AT1G15540.1 | Fe(II)-dependent oxygenase superfamily protein |
| 1 | Medtr1g100787 | 45717233 | 1.75E-08 | 0.14 | 15.2 | intergenic | AT5G07280.1 | Leucine-rich repeat transmembrane protein kinase |
| 2 | Medtr2g438720 | 15656689 | 5.61E-08 | 0.13 | 13.8 | non-synonymous |  | Ankyrin repeat plant-like protein |
| 8 | Medtr8g015980 | 5289479 | 4.56E-07 | 0.11 | 15.0 | non-synonymous | AT3G21250.2 | ABCC8; transmembrane ATPase activity |
| 5 | Medtr5g070320 | 29773843 | 6.99E-07 | 0.10 | 10.2 | intron | AT2G29940.1 | PDR3; pleiotropic drug resistance 3 |
| 5 | Medtr5g070330 | 29783407 | 1.96E-06 | 0.10 | 9.3 | intron | AT3G51860.1 | CAX3; cation exchanger 3 |
| 2 | Medtr2g069090 | 28697944 | 2.08E-06 | 0.10 | 12.2 | downstream | AT5G43440.1 | Fe(II)-dependent oxygenase superfamily protein |
| 5 | Medtr5g070330 | 29782526 | 3.26E-06 | 0.09 | 8.7 | 3 prime UTR | AT2G29940.1 | PDR3; pleiotropic drug resistance 3 |
| 3 | Medtr3g090170 | 40983523 | 4.49E-06 | 0.09 | 9.3 | intron | AT1G77140.1 | ATVPS45; vacuolar protein sorting 45 |
| 2 | Medtr2g019020 | 6102461 | 7.32E-06 | 0.09 | 10.1 | synonymous | AT2G34660.2 | ABCC2; transmembrane ATPase activity |
| 2 | Medtr2g095480 | 40788857 | 9.99E-06 | 0.09 | 12.7 | upstream | AT5G60800.2 | HIPP3; Heavy metal transport/detoxification superfamily protein |

Hg leaf
| Chr | Gene Name | Position | p-value | PVE | Effect Size | location | BLAST *A. thaliana* | Gene Description |
| --- | --- | --- | --- | --- | --- | --- | --- | --- |
| 3 | Medtr3g088460 | 40127485 | 1.15E-07 | 0.14 | 136.5 | intron | AT1G15960.1 | NRAMP6; metal ion transporter 6 |
| 3 | Medtr3te114660 | 53546134 | 4.95E-07 | 0.13 | 117.4 | downstream | AT1G20270.1 | Fe(II)-dependent oxygenase superfamily protein |
| 2 | Medtr2g036380 | 15751342 | 8.27E-06 | 0.10 | 106.6 | intron | AT4G30110.1 | HMA2; heavy metal atpase 2 |
| 5 | Medtr5g033320 | 14374444 | 1.96E-05 | 0.09 | 114.2 | intron | AT3G13080.1 | ABCC3; transmembrane ATPase activity |

Hg RRG
| Chr | Gene Name | Position | p-value | PVE | Effect Size | location | BLAST *A. thaliana* | Gene Description |
| --- | --- | --- | --- | --- | --- | --- | --- | --- |
| 2 | Medtr2g083420 | 35018740 | 8.10E-10 | 0.18 | 16.5 | non-synonymous | AT4G09500.2 | UDP-Glycosyltransferase superfamily protein |
| 4 | Medtr4g116010 | 47960277 | 2.51E-06 | 0.10 | 16.2 | intergenic | AT1G49390.1 | Fe(II)-dependent oxygenase superfamily protein |
| 2 | Medtr2te008720 | 1586136 | 3.51E-06 | 0.10 | 13.0 | downstream | AT2G41480.1 | Peroxidase superfamily protein |
| 5 | Medtr5g073020 | 31072945 | 6.78E-06 | 0.09 | 12.7 | intron | AT3G26210.1 | Cytochrome P450, family 71, subfamily B, polypeptide 23 |
| 3 | Medtr3g014705 | 4234570 | 1.27E-05 | 0.09 | 16.6 | intron | AT3G51950.2 | Zinc finger (CCCH-type) family protein / RNA recognition motif |
| 1 | Medtr1g063920 | 28084622 | 1.66E-05 | 0.08 | 8.8 | intron | AT2G37010.1 | non-intrinsic ABC protein 12 |
| 1 | Medtr1g084780 | 37746862 | 1.47E-05 | 0.08 | 17.2 | downstream | AT4G30190.2 | H(+)-ATPase 2 |
| 4 | Medtr4g076900 | 29426248 | 1.67E-05 | 0.08 | 16.3 | intron | AT1G51500.1 | ABC-2 type transporter family protein |

Using GWAS and population genomic scans (F_st_, and XP-CLR), we found a large region (∼1.3 Mb) of interest associated with Cd RRG on chromosome 2. A highly significant non-synonymous SNP was found at position 15,656,689 (p-value = 5.61 × 10^−8^) in an ankyrin repeat protein (Medtr2g438720), and the surrounding region contained multiple ankyrin repeat genes within 1 kb of many top SNPs (**Figure 4A, Table 2**). Eleven of those SNPs were in exons, nine of which were non-synonymous SNPs, while two others were synonymous (**Supplementary Table 5**). The high LD in this region suggested the ankyrin genes were linked. While these ankyrin genes did not show homology to *A. thaliana* orthologs, some ankryin repeat proteins have been shown to mediate membrane bound protein-protein interactions that facilitate heavy metal transport in *A. thaliana* roots (Gao et al., 2009). Genomic divergence between high and low Cd RRG genotypes was found by scanning for high F_st_ in a region upstream from the GWAS peak (**Figure 4B**). This region contains a tandemly triplicated ABC transporter (*MtABCC14*, Medtr2g436680; **Supplementary Figure 5**), an ortholog of Heavy Metal Transporter 10 (*MtHMP10*, Medtr2g436830), a Fe (II)-dependent oxygenase superfamily protein, ATPase transporter *MtHMA7* (Medtr2g035840), and a potassium transporter (Medtr2g438160; **Table 2**). A significant peak was also detected using XP-CLR for the window containing the three *MtABCC14* copies and *MtHMP10* (Medtr2g436830; **Figure 4C, Table 2**), which was the same genomic region identified using F_st_.

**Figure 4.**
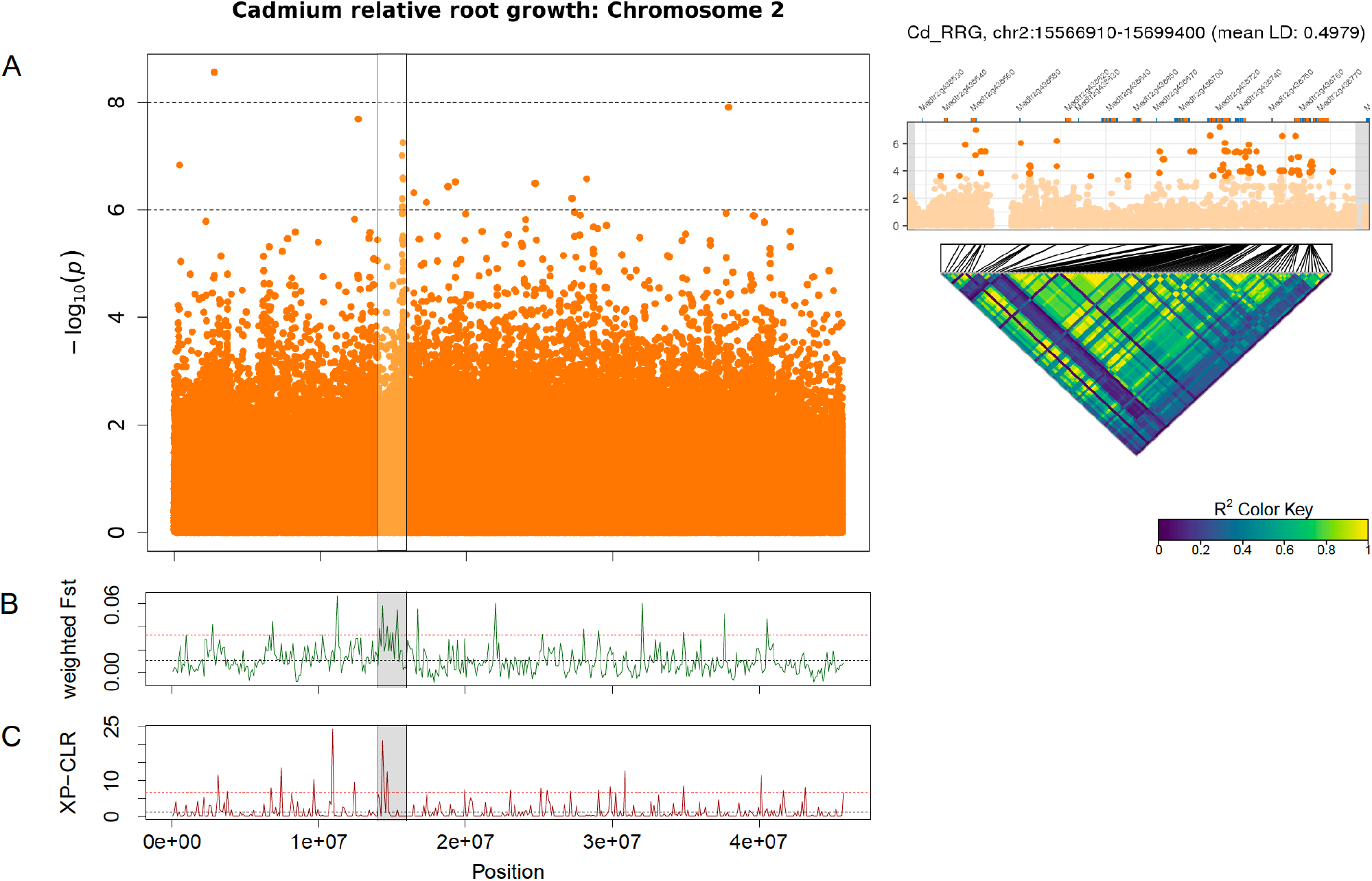
Manhattan plot of Cd relative root growth (RRG) associations on chromosome 2 using GAPIT (A). The highlighted cluster of SNP’s (horizontal black bars) contains multiple ankyrin repeat genes (see Table 2). Each point represents a SNP, the x-axis signifies the position on the chromosome while the y-axis is the -log_10_ of the p-value. Pairwise linkage disequilibrium (LD) among the 100 most significant SNPs in the peak is shown as a heatmap on the right, the most significant SNPs and Mt4.0 genes in this region shown above the heatmap. The black lines mark the position of each SNP in the peak, dark blue color signifies no LD, green to yellow is high to maximum LD. Weighted F_st_ statistics from sliding window analysis (100 kb windows) on chromosome 2 with GWAS peak marked in grey (B). Higher F_st_ indicates genomic differentiation based on the groups of plant genotypes with the highest vs. lowest Cd RRG values. XP-CLR is a two-group comparison of selective sweeps using plant genotypes with the highest vs. lowest Cd RRG values (C). The dashed black line represents the mean F_st_ or CLR values for the chromosome, the dashed red line represents windows ≥ 5% of the highest F_st_ or CLR values for the chromosome.

**Table 2.** Genes in chromosome 2 genomic regions associated with Cd relative root growth (Cd RRG) that were identified using GWAS, F_st_ and XP-CLR. Columns from left to right are GO-terms, Mt4.0 gene ID, *A. thaliana* ortholog gene function identified by BLAST, and gene location with start and stop coordinates based on the Mt4.0 genome.

**GWAS**
| GO-term | Mt4.0 gene | BLAST gene function | Mt4.0 location |
| --- | --- | --- | --- |
| integral component of membrane | Medtr2g438720 | ankyrin repeat plant-like protein | chr2:15654113..15660513 |
| integral component of membrane | Medtr2g438740 | ankyrin repeat plant-like protein | chr2:15661862..15664982 |
| integral component of membrane | Medtr2g438760 | ankyrin repeat plant-like protein | chr2:15678476..15683145 |
| integral component of membrane | Medtr2g438700 | ankyrin repeat plant-like protein | chr2:15644925..15649215 |
|  | Medtr2g438560 | hypothetical protein | chr2:15587523..15589330 |
| integral component of membrane | Medtr2g438580 | ankyrin repeat protein | chr2:15601177..15601431 |
|  | Medtr2g438670 | hypothetical protein | chr2:15639667..15639882 |

**$F_{st}$ (5% tail)**
| GO-term | Mt4.0 gene | BLAST gene function | Mt4.0 location |
| --- | --- | --- | --- |
| ATPase activity, transmembrane movement | Medtr2g436680 | ABC transporter protein (ABCC14, AT3G62700.1) | chr2:14273804..14283871 |
| ATPase activity, transmembrane movement | Medtr2g436710 | ABC transporter protein (ABCC14, AT3G62700.1) | chr2:14295748..14305736 |
| ATPase activity, transmembrane movement | Medtr2g436730 | ABC transporter protein (ABCC14, AT3G62700.1) | chr2:14312635..14321998 |
| metal ion binding/ transport | Medtr2g436830 | heavy metal transport (ATHMP10, AT1G56210.1) | chr2:14380972..14382489 |
| metal ion binding | Medtr2g437380 | 2OG-Fe(II) oxygenase family oxidoreductase | chr2:14380972..14382489 |
| ATPase activity, metal ion binding/transport | Medtr2g035840 | heavy metal P-type ATPase (HMA7, AT5G44790.1) | chr2:15206300..15212503 |
| potassium ion transmembrane transport | Medtr2g438160 | potassium transporter 12 (AT1G31120.1) | chr2:15394059..15398083 |

**XP-CLR (5% tail)**
| GO-term | Mt4.0 gene | BLAST gene function | Mt4.0 location |
| --- | --- | --- | --- |
| ATPase activity, transmembrane movement | Medtr2g436680 | ABC transporter protein (ABCC14, AT3G62700.1) | chr2:14273804..14283871 |
| ATPase activity, transmembrane movement | Medtr2g436710 | ABC transporter protein (ABCC14, AT3G62700.1) | chr2:14295748..14305736 |
| ATPase activity, transmembrane movement | Medtr2g436730 | ABC transporter protein (ABCC14, AT3G62700.1) | chr2:14312635..14321998 |
| metal ion binding/ transport | Medtr2g436830 | heavy metal transport (ATHMP10, AT1G56210.1) | chr2:14380972..14382489 |
| metal ion binding | Medtr2g437380 | 2OG-Fe(II) oxygenase family oxidoreductase | chr2:14380972..14382489 |

Another cluster of low p-values associated with Cd RRG was found on chromosome 5 (**Supplementary Table 6**) with SNPs in high LD (**Figure 5A**). When we ran the GWAS following log_2_ transformation of the Cd RRG phenotype, the model fit was somewhat improved, and this peak became more pronounced (**Figure 2, Supplementary Figure 4**). In this genomic region, five SNPs (four intronic SNPs and one in the 3’-UTR) were in *MtCAX3* (Cation Exchanger 3, Medtr5g070330), and three SNPs in the Pleiotropic Drug Resistance 3 gene, *MtPDR3* (Medtr5g070320). In the Mt4.0 reference genome, *MtCAX3* and *MtPDR3* are adjacent (5878 bp apart; **Supplementary Figure 6**). The peak also contained a non-synonymous, synonymous, and several intronic SNPs in a damaged DNA-binding protein, *MtDDB2* (Medtr5g070310), and an undecaprenyl pyrophosphate synthase *MtCPT7* (Medtr5g070270), which is orthologous to *AtCPT7* (also known as *AtCPT4*). We found several intergenic SNPs with low p-values in high LD in the region containing a cluster of duplicated undecaprenyl pyrophosphate synthase genes (Medtr5g070210, Medtr5g070220, Medtr5g070230; **Figure 5**) that are upstream from *MtCPT7*, all of which appear to be linked. The window containing *MtCAX3* and *MtPRD3* is also in the upper tail of F_st_ on chromosome 5 (**Figure 5B**), and the XP-CLR statistic showed a significant peak just upstream from the GWAS peak (**Figure 5C**).

**Figure 5.**
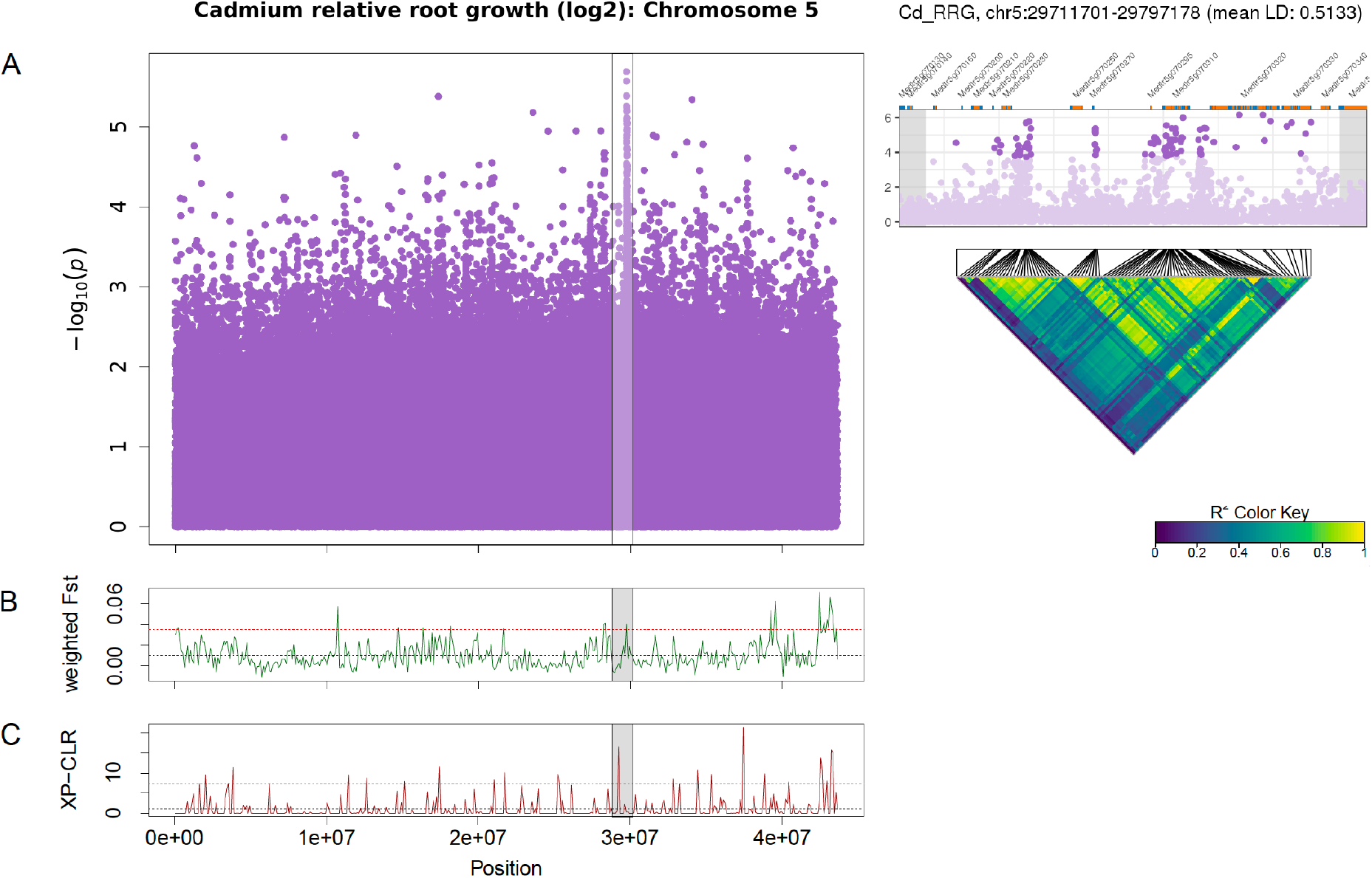
Manhattan plot of Cd relative root growth (RRG) associations for chromosome 5 using GAPIT (A). The region on chromosome 5 associated with Cd RRG contained *MtCAX3* (Medtr5g070330), *MtPDR3* (Medtr5g070320), *MtCPT7* (Medtr5g070270) and *MtDDB2* (Medtr5g070310). The SNPs in genic and intergenic regions are in high LD (pairwise linkage disequilibrium (LD) among the 100 most significant SNPs in the peak). Each point represents a SNP, the x-axis signifies the position on the chromosome while the y-axis is the -log_10_ of the p-value. The Cd RRG phenotype was log_2_ transformed to normalize the data prior to running the GWAS. The GWAS region is highlighted in gray the F_st_ (B) and XP-CLR (C) sliding window (100 kb windows) analyses. The x-axis represents the chromosomal position, the y-axis the F_st_ or XP-CLR value of the window at the corresponding position. Higher F_st_ indicates population differentiation, higher XP-CLR the presence of a selective sweep. The dashed black line represents the mean F_st_ or CLR values for the chromosome, the dashed red line represents windows ≥ 5% of the highest F_st_ or CLR values for the chromosome.

For Hg leaf accumulation, one of the most significant SNPs was found in the Natural Resistance Associated Macrophage Protein 6 gene (*MtNRAMP6*, Medtr3g088460). In the *M. truncatula* A17 genotype, *MtNRAMP6* is expressed most highly in the roots (**Supplementary Dataset 1**). Genes found among the top 1000 SNPs were the heavy-metal ATPase *MtHMA2* (Medtr2g036380) and *MtABCC3* (Medtr5g033320). We found a large region (∼2.3 Mb) on chromosome 8 (**Figure 6**) that contains a non-synonymous SNP in *MtCAX9* (Medtr8g085260) and several surrounding genes with ATP binding/ATPases, and transmembrane GO-terms (**Supplementary Table 7**). The top 100 SNPs in this region showed very high LD (*r*^2^ = 0.62), suggesting a large locus with several linked genes that may be involved in Hg accumulation. This region overlaps with high F_st_ windows spanning the same chromosomal region, which indicates higher Hg accumulating genotypes are genetically differentiated in this large genomic region. Among the genes in the highest F_st_ peak included a duplicated cation amino acid transporter (four homologs: Medtr8g089320, Medtr8g089340, Medtr8g089342, Medtr8g089360) a magnesium-translocating P-type ATPase (Medtr8g089870), an ATPase amine-terminal autoinhibitory domain protein (Medtr8g090120) and an adjacent membrane calcium-translocating P-type ATPase (Medtr8g090125) (**Supplementary Table 8**). The population genomics methods provided complementary support that the regions on chromosomes 2, 5, and 8 identified by GWAS contain a genomic signal that could be detected using only the genotypes with the most extreme phenotypes.

**Figure 6.**
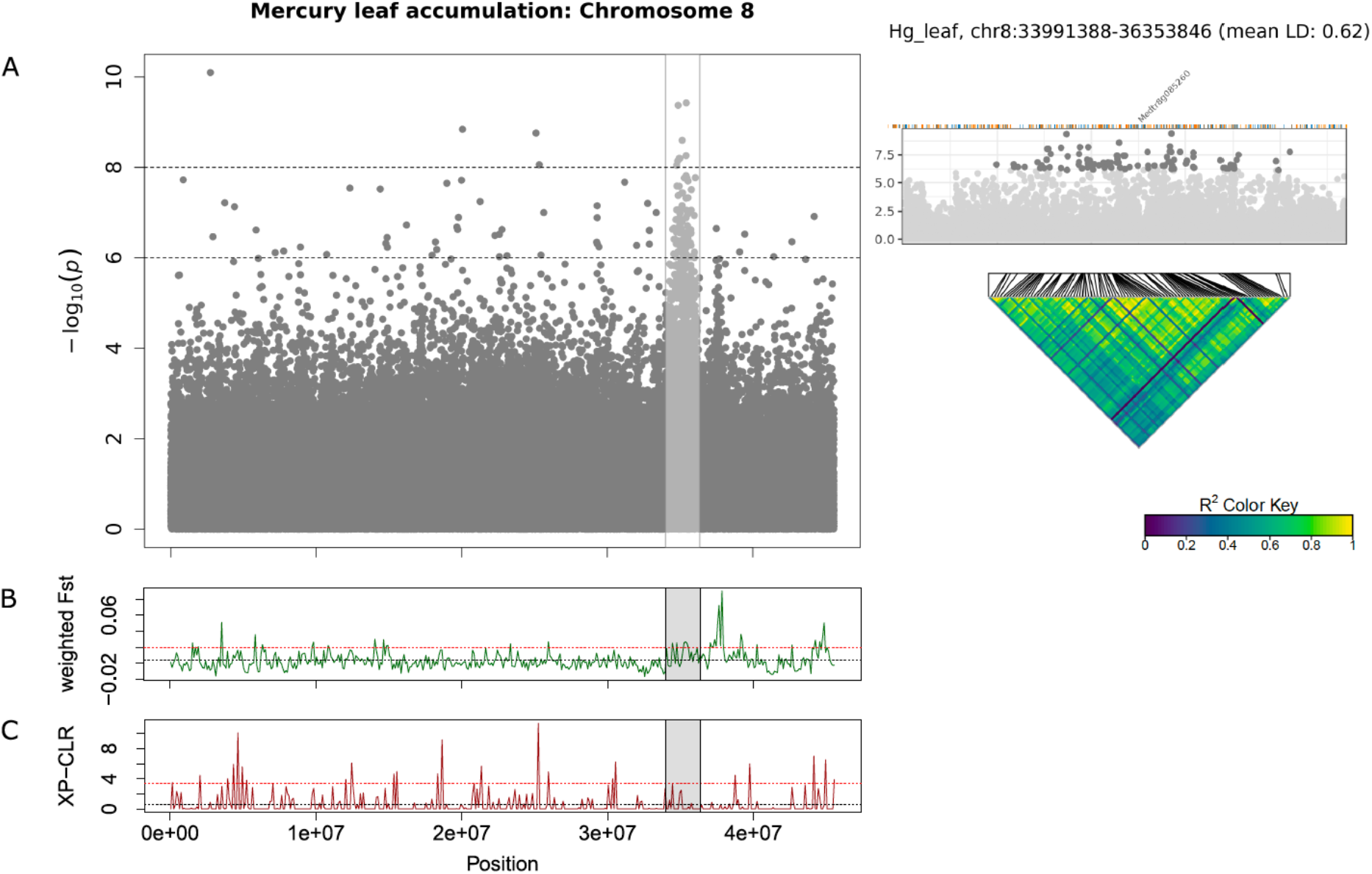
Manhattan plot of Hg leaf accumulation for chromosome 8 using GEMMA (A). Each point represents a SNP, the x-axis signifies the position on the chromosome while the y-axis is the -log_10_ of the p-value. The Cd RRG phenotype was log_2_ transformed to normalize the data prior to running the GWAS. The GWAS region is highlighted in gray the F_st_ (B) and XP-CLR (C) sliding window (100 kb windows) analyses. The x-axis represents the chromosomal position, the y-axis the F_st_ or XP-CLR value of the window at the corresponding position. Higher F_st_ indicates population differentiation, higher XP-CLR the presence of a selective sweep. The dashed black line represents the mean F_st_ or CLR values for the chromosome, the dashed red line represents windows ≥ 5% of the highest F_st_ or CLR values for the chromosome.

For Hg RRG, the top SNP was a non-synonymous substitution on chromosome 2 in a UDP-glycosyltransferase gene (Medtr2g083420). This gene was shown to be most highly expressed in roots in the A17 genotype (**Supplementary Dataset 1**). Among the other top SNPs for Hg RRG was a Fe (II)-dependent oxygenase superfamily protein (Medtr4g116010) and a peroxidase superfamily protein, both with metal ion binding and oxidoreductase activity GO-terms, and a cytochrome P450 gene (Medtr5g073020) with heme binding, membrane, and metal ion GO-terms. Three ATP binding genes all of which have membrane/transport ontologies, including H^+^ ATPase 2 (Medtr1g084780), non-intrinsic ABC protein 12 (Medtr1g063920), and an ABC-2 type transporter family protein (Medtr4g076900) (**Table 1**), were also associated with Hg RRG.

### Minor Allele Frequency and Effect Size

The distribution of effect sizes of all SNPs is centered around zero, while those with the lowest p-values are shifted significantly toward larger effect sizes (**Supplementary Figure 7**). For all four trait associations, we found negative correlations between MAF’s and effect sizes (**Figure 7**), where SNPs with larger effect sizes had the tendency to be at low frequency. This is consistent with purifying selection acting to remove alleles that may be deleterious if they result in elevated toxicity after exposure to Cd or Hg. Conversely, SNPs with large effect size associated with lower leaf accumulation or lower RRG typically had higher MAF’s, which may be consistent with positive selection for alleles that reduce uptake of toxic metals. Because SNPs with the lowest p-values are inherently biased towards large effect sizes, we ran GWAS on 100 permuted phenotype datasets to generate a null distribution for each trait. For Cd leaf accumulation and RRG, the correlation coefficients from the empirical datasets were below 99 and 86 of the 100 permuted datasets, respectively (**Supplementary Figure 8**). For Hg leaf accumulation and RRG, the correlation coefficients from the empirical dataset coefficients were below only 37 and 52 of the permuted datasets. Among the four traits, we would only consider the correlation coefficient from Cd leaf accumulation to be statistically significant (p = 0.01). Nevertheless, the correlations between MAF and effect size in all four traits were consistently negative. We used the Tajima’s D statistic to test for significant differences in the top SNPs detected in GWAS compared with the rest of the genome (i.e., “background SNPs”). We found significantly lower mean Tajima’s D for the top SNPs from the GWAS compared with background SNPs for three of the four traits (**Figure 8**) due to an excess of low frequency alleles (i.e., low MAF’s), consistent with purifying selection (Paape et al., 2013).

**Figure 7.**
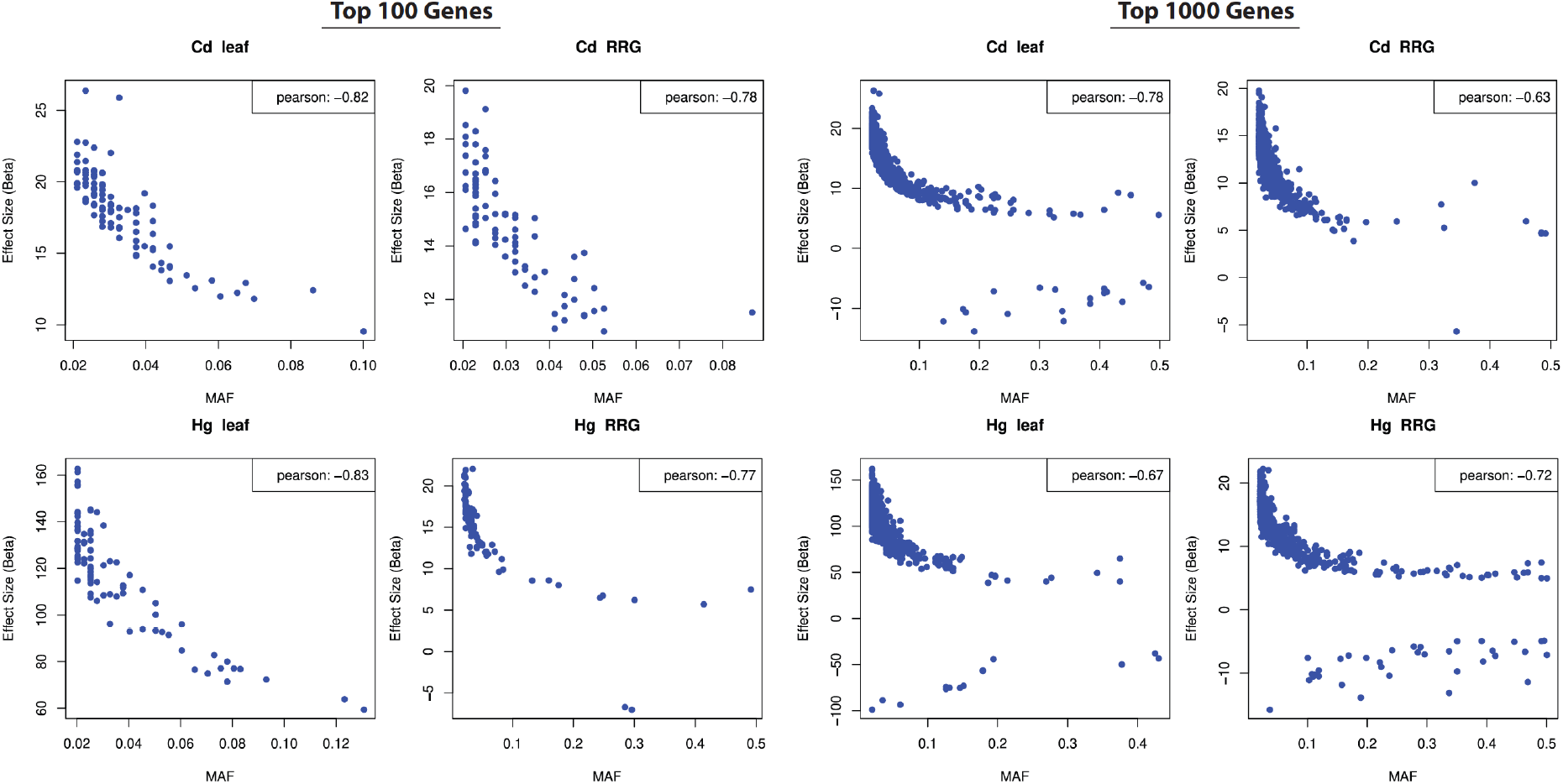
Correlations between minor allele frequency (MAF; x-axis) and SNP effect size (y-axis) in the top 100 (left four panels) and top 1000 (right four panels) most significant SNPs (effect sizes estimated from GAPIT). Pearson’s correlation coefficients shown in upper right corners.

**Figure 8.**
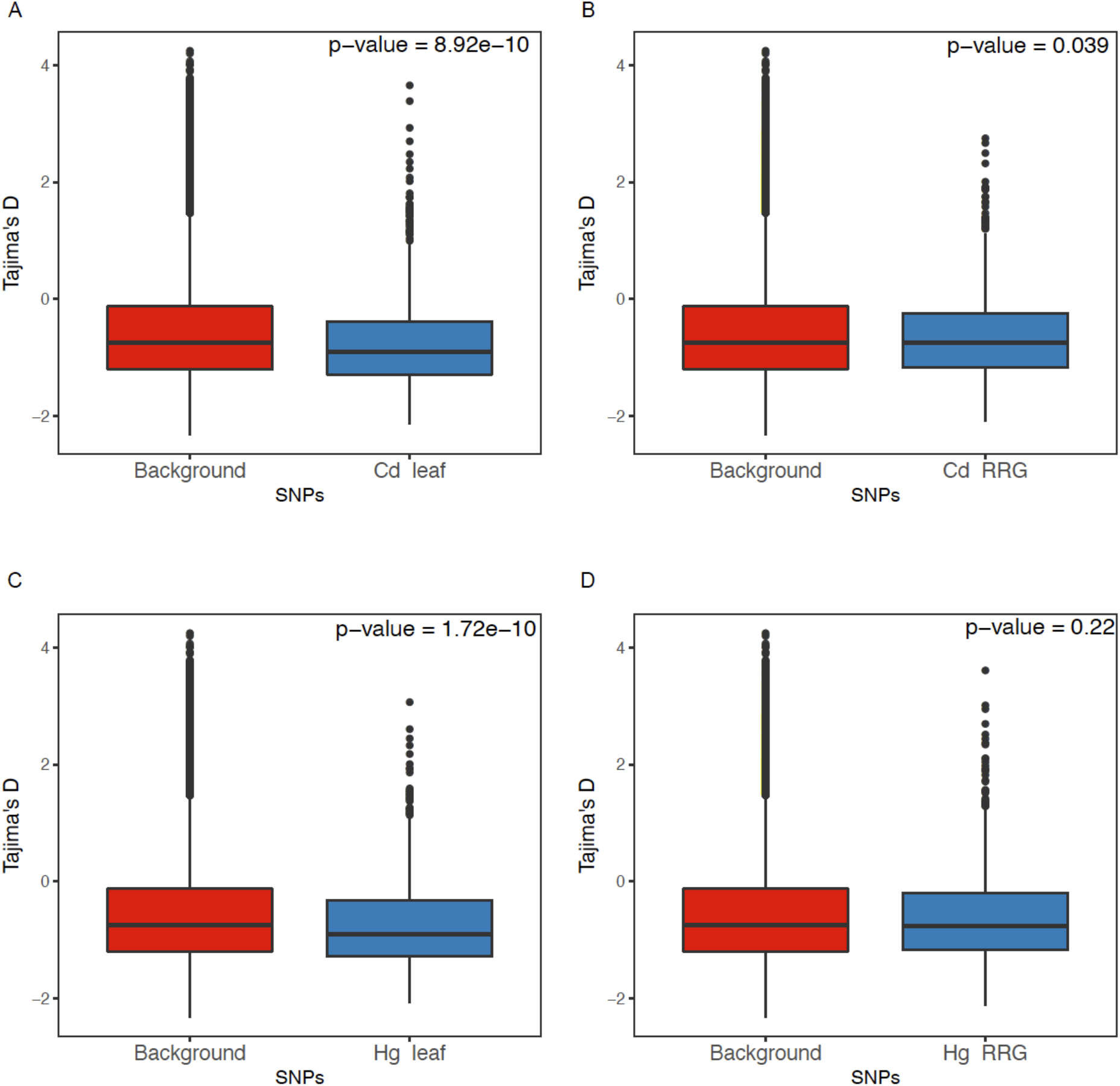
Comparison of Tajima’s D from the genomic background and regions containing SNPs from the GWAS for each of the traits. Tajima’s D was estimated using 10 bp regions surrounding focal SNPs.

## Discussion

### Phenotypic and Genetic Diversity

Heritability was sufficiently high for GWAS and our heritability estimates showed similar levels of phenotypic/genetic variability for the two RRG traits (about 10% difference) as well as for the leaf accumulation traits. Overall, heritability was higher for RRG than for leaf accumulation. Variability in Cd accumulation found in *M. truncatula* leaves was comparable to findings in *A. thaliana* (Chao et al., 2012), barley (Wu et al., 2015), *Brassica napus* (Chen et al., 2018), and rice (Zhao et al., 2018). Interestingly, variation in Hg accumulation in leaves was about 5 times wider than Cd accumulation in *M. truncatula*. To our knowledge, the only other study that has measured Hg accumulation in a wild plant or crop association panel was in maize (Zhao et al., 2017). Our study showed significantly higher variation in leaf Hg accumulation than the maize study (∼270-fold versus ∼3.5-fold). However, Zhao et al. (2017) collected mature plant leaves under significantly different conditions, making direct comparisons between studies difficult. Our phenotyping approach also revealed no correlation between RRG and leaf accumulation in *M. truncatula*, indicating that high metal tolerance does not translate to higher metal accumulation in leaves (see Figure 1). The strongest correlation was between the RRG of plants treated with Cd and Hg. Our findings are similar to a recent study comparing RRG responses to Cd and Hg treatments in a different collection of *M. truncatula* genotypes (*r* = 0.48) which did not include sequence data (García de la Torre et al., 2021). The positive correlation in RRG between the two metal treatments suggests that both heavy metals elicit a common stress response that inhibits root growth while retaining other metal-specific responses (i.e., the correlation, *r* < 0.5), but existing genetic diversity for root length may also be a contributing factor.

### Candidate Genes are Broadly Dispersed Across the Genome

In many cases the most significant SNPs were found in functionally relevant genes that were broadly dispersed throughout the genome. For this reason, we found it more meaningful to examine a wide range of p-values (i.e., the top 100 or top 1000) for each trait association because we expected RRG and leaf accumulation of Cd and Hg to be highly polygenic. This allowed us to look at gene annotations and biologically relevant GO-terms among a list of several hundred genes for each trait. Our findings for Cd accumulation are considerably different than a similar GWAS in *A. thaliana*, where a single genomic peak containing the *HMA3* locus was identified, which drowned out the signal at any other genomic region (Chao et al., 2012). We found clusters of SNPs in multiple genomic regions, all of which appeared to have several linked genes with functional relevance to metal transport or metal tolerance. In cases such as on chromosomes 2, 5, and 8, the genomic region of interest appears to constitute a large locus containing many functionally relevant genes. If these genes are functionally linked, they likely evolved together as a single locus, such as *MtCAX3* and *MtPDR3* on chromosome 5 which are clearly linked. Moreover, the detection of orthologs that have functionally relevant annotations indicates that our GWAS provides a realistic picture of the genetic architecture underlying metal tolerance traits in *M. truncatula*. Included among the genes identified in our genome wide statistical analyses were several stress-related genes, metal ion transporters, membrane transporters, and vacuole proteins with ion binding gene ontologies. Many of these genes are orthologous to known metal transporters studied in other plant species suggesting they are evolutionarily conserved.

A large category of candidate genes were ATP-dependent transporters that were associated with all traits, suggesting that ATP binding genes are important for Cd and Hg tolerance in *M. truncatula*. Included among these were multiple ATP binding cassette (ABC) proteins associated with each of the traits. Many of these are orthologous to ABC-transporters in *A. thaliana* (e.g., *AtABCC2, AtABCC3, AtABCC8*) and rice that are known to interact with metal-binding phytochelatins that chelate Cd and Hg ions allowing for transportation into vacuoles by membrane bound ABC-proteins (Park et al., 2012; Brunetti et al., 2015; Sun et al., 2018). Another class of ATP-transporters are the P_1B_-type heavy metal ATPases (HMA’s) which are well-known for their roles in root to shoot transport (Hanikenne et al., 2008) and compartmentalization of unchelated heavy metals (Morel et al., 2008). HMA’s have been shown to play essential roles in Cd, Co, Pb and Zn transport in *A. thaliana* and rice (Chao et al., 2012; Takahashi et al., 2012). Thus far, HMAs have not been associated with Hg tolerance and transport in plants, instead relying on ABC-transporters (Park et al., 2012). In our study, we found *MtHMA2* associated with leaf accumulation of Hg and a P_1B_-type heavy metal ATPase (H^+^-ATPase 2) associated with Hg RRG. Interestingly, no HMA’s were associated with either of the Cd traits. Clearly, ABC- and ATPase transporters have widespread roles that range from general to specific metal responses in plants, which include transport of essential and non-essential metal ions (Takahashi et al., 2012).

The most strongly associated candidate gene for Hg leaf accumulation was the Natural Resistance-Associated Macrophage Protein 6, *MtNRAMP6*. In rice and *A. thaliana, NRAMP’s* are involved in aluminum (Xia et al., 2010) and manganese transport (Cailliatte et al., 2010). This gene has the highest expression in roots which may result in greater availability of Hg to be transported to the leaf leading to higher concentrations in the high Hg-accumulating genotypes. In *A. thaliana*, overexpressed *NRAMP6* resulted in Cd hypersensitivity through increased Cd uptake, while *nramp6* loss of function mutants resulted in reduced Cd uptake leading to increased plant tolerance when exposed to high levels of Cd (Cailliatte et al., 2009). Similarly, in rice, knock-out mutations of *NRAMP5* resulted in reduced Cd accumulation in roots and leaves (Sasaki et al., 2012). Previous GWAS studies in barley and rapeseed found that homologs of *NRAMP5* and *NRAMP6* were associated with Cd accumulation (Wu et al., 2015; Chen et al., 2018). While our study found an association between *NRAMP6* and Hg, but not Cd accumulation, our results provide supporting evidence that this family of genes contributes to tolerance and transport of an array of metal ions using similar mechanisms (Nevo and Nelson, 2006).

### Large Genomic Regions Identified with Multiple Methods

We also found genomic regions with clusters of SNPs and genes that showed linkage for Cd tolerance and leaf accumulation of Hg, which aligned with regions also identified using population genomics methods. Using genotypes that were divergent for Cd RRG, the F_st_, and XP-CLR statistics identified a region on chromosome 2 that containing a tandemly triplicated ABC-transporter (*MtABCC14*), the heavy metal transport/detoxification superfamily protein *AtHMP10*, and a Fe-oxidoreductase gene. Interestingly, this region has very high LD and contains several gene families with known metal tolerance roles and gene ontologies that support metal transport mechanisms or metal-induced stress responses. Expression enhancement through the triplication of genes is a common mechanism in metal ion transport in hyperaccumulating plants (Hanikenne et al., 2008; Shahzad et al., 2010) suggesting that the three linked *MtABCC14* copies may jointly contribute to tolerance to Cd. A second F_st_ peak containing the P_1B_-type heavy metal ATPase *MtHMA7* and a potassium transporter was found closer to the GWAS peak that contained ankyrin repeats. While these ankyrin genes do not show homology to *A. thaliana* orthologs, ankryin repeat proteins were shown to mediate membrane bound protein-protein interactions that facilitated heavy metal transport in *A. thaliana* roots (Gao et al., 2009). In a GWAS study on salinity tolerance in *M. truncatula*, chromosome 2 was previously shown to have many loci associated with ion stress, with the closest peak being approximately 800 kb away from the peak identified here (Kang et al., 2019). We propose that this arm of chromosome 2 has clusters of specific and general ion transporters that influence both toxic and essential ions in *M. truncatula*.

The region on chromosome 5 associated with Cd RRG was identified using both GWAS and F_st_ statistics, and contains the genes *MtCAX3* and *MtPDR3* which are located < 6 kb apart. Homologs of these two genes have been shown to have substantial roles in heavy metal tolerance. The cation exchanger genes *CAX1* and *CAX3* are transporters located on the tonoplast. Upregulation of *CAX3* was reported to increase Cd tolerance in *A. thaliana* by sequestering Cd into the vacuole (Cheng et al., 2005). Expression levels of *CAX3* are upregulated by Hb1 (class 1 hemoglobin), which was shown to inhibit the expression of *IRT1* and *PDR8* (Bahmani et al., 2019). The ATP-binding cassette transmembrane transporter *PDR3* is an ortholog of *PDR8*. Inhibition of *PDR8* using RNAi was shown to increase sensitivity to Cd and overexpression increased resistance to Cd in *A. thaliana* (Kim et al., 2007). Because *PDR3* and *PDR8* are both ABC-transmembrane proteins, we assume their function is similar, as we have seen for other ABC-proteins involved in heavy metal transport to vacuoles. Another gene that is near the linked *MtCAX3* and *MtPDR3* genes is *MtCPT7*, which is homologous to *AtCPT7* (also known as *AtCPT4*). *AtCPT7* is involved in dolichol synthesis and was shown to be upregulated after exposure to Cd in *A. thaliana* (Jozwiak et al., 2017). Because *MtCAX3, MtPDR3*, and *MtCPT7* have biologically plausible roles in Cd transport, their tight linkage suggests they may be collectively involved in Cd tolerance.

To date, few QTL associated with Hg accumulation have been identified in crops or crop relatives (but see Wang et al., 2013; Zhao et al., 2017). Using GWAS, we found a large genomic region of interest on chromosome 8 in *M. truncatula* that had several transmembrane and ATPase transporters in a region surrounding *MtCAX9*. As discussed above, the *CAX* transporters have been found to be located on vacuole membranes, and the large number of membrane proteins and transmembrane related gene ontologies (24 genes spanning ∼1.2 Mb) suggests this genomic region possesses biologically relevant candidate genes involved in transport of Hg across membranes, including into vacuoles. On this chromosome, the highest F_st_ peak that is upstream from the GWAS peak includes multiple calcium and magnesium related P-type ATPases. This finding fits the recurring theme that P-type ATPases are involved in multiple metal ion transport mechanisms, including for highly toxic ions, such as Hg. Chromosome 8 contains some highly promising candidate genes to pursue using functional genetics approaches such as knock-out mutations using Hg accumulation to quantify the effects of mutants.

### Stress Related Genetic Responses

Together, cellular responses that increase tolerance or accumulation to heavy metals are complex and involve not only ion transporters, but also mechanisms that deal with toxicity such as oxidative stress and ROS production. In excess amounts, all heavy metals trigger oxidative stress responses, resulting in increased levels of ROS in cells and tissues. As storage capacity is exceeded and toxic ions reach the leaves, plants alter leaf structure and chemistry leading to an accumulation of hydroxyl radicals. These hydroxyl radicals will indiscriminately oxidize biomolecules, including vital proteins and plant enzymes. In *M. truncatula*, Cd tolerance has been associated with the efficiency of the plant’s antioxidant defenses (Rahoui et al., 2014). The strong association with multiple Fe-oxidoreductases with all four traits suggests that plant tissues are likely under oxidative stress in our study. Moreover, *MtOSX2*, which has metal ion binding gene ontology, is associated with Cd leaf accumulation and may also play a role in dealing with oxidative stress responses to heavy metals (Kawai et al., 2014). In *A. thaliana*, the *AtOXS2* gene (also known as *DEG9*) is a zinc-finger transcription factor shown to be triggered by salt stress and in maize (*Zea mays*), *ZmOXS2* homologs are able to enhance Cd tolerance (He et al., 2016; Jing et al., 2019). The most strongly associated SNP for Hg RRG was a non-synonymous substitution in a UDP-glycosyltransferase gene (Medtr2g083420), which is known to have highest expression in the roots of the A17 ecotype. UDP-glycotransferase mutants of *AtUGGT* in *A. thaliana* were shown to have delayed root growth induced by ion stress, which involved disruption of endoplasmic reticulum transmembrane integrity (Blanco-Herrera et al., 2015). In *M. truncatula*, UDP-glycosyltransferases are involved in the biosynthesis of plant secondary metabolites, including flavonoids and isoflavonoids (Modolo et al., 2007). Flavonoids are crucial anti-oxidants that can efficiently scavenge the harmful ROS generated by heavy metal toxicity. They have also been reported to act as metal chelators (Kumar and Prasad, 2018).

### Standing Variation and Genetic Architecture

The top SNPs detected in our GWAS showed a strong negative relationship between MAF and effect size, consistent with expectations of mutation-selection balance (Barton and Keightley, 2002) and stabilizing selection (Josephs et al., 2017). While some mutations may result in alleles that confer greater accumulation or tolerance to toxic heavy metals that could be beneficial, we expect that most mutations that result in excess uptake of heavy metals to be deleterious. This explains why alleles with large effect that were associated with high tolerance or accumulation were at low frequencies in the population/species because we expect purifying selection to act on these mutations. The SNPs with large effect that are associated with lower high tolerance or accumulation but are at higher frequencies, would be expected to be more beneficial because they resulted in exclusion of Cd or Hg, and would therefore be more common in the species. However, a large amount of standing genetic variation, including SNPs associated with higher tolerance or accumulation, may provide the necessary polymorphism for species to colonize or adapt to new habitats (Hermisson, 2005; Barrett and Schluter, 2008), suggesting potential benefits of these alleles in some environments. Hence, trade-offs between beneficial and deleterious mutations are expected under models of stabilizing or balancing selection which favors intermediate phenotypes and allele frequencies. It is also likely that alleles conferring greater tolerance or accumulation of Cd or Hg have been tested in natural or contaminated environments at some point throughout the species history and may have enabled a broader species distribution. Moreover, many alleles found in our GWAS to be associated with these specific heavy metals, may be generally associated with stressful levels of ion contamination due to poor quality soils and not necessarily due to direct heavy metal contamination from mining in the very recent past.

## Conclusion

By combining GWAS analysis with population genomics, we were able to find signals of trait associations that overlapped with genomic regions detected by GWAS. Our population genomics approach used the tails of quantitative trait distributions to provide a contrast for genome-wide scans of genomic divergence, similar to the QTL approach described by Stinchcombe and Hoekstra (2007), but using the natural variation in the Medicago HapMap panel. With this in mind, it could be feasible to sequence only those lines in the ends of the phenotypic distributions in very large collections of genotypes (e.g., García de la Torre et al., 2013, 2021), and find meaningful genetic loci underlying the trait differences without sequencing an entire GWAS panel. This approach likely depends on phenotyping a rather large collection of individual genotypes to ensure that the ends of the phenotypic distributions are truly representative of the species level ranges in trait values. Genome-wide association mapping and population genomic scans revealed that leaf and root responses to toxic heavy metals are complex polygenic traits involving many quantitative trait loci. We found ABC-transporters and P-type ATPases to be commonly associated with Cd and Hg tolerance, which is supported by previous functional studies among members of these gene families playing large roles in heavy metal transport across membranes and isolation in vacuoles. While genetic architect may vary widely across plant species to due to the spatial and genomic distribution of genes involved in metal ion transport, accumulation, and detoxification, gene function of the families of transporters and stress response genes seems to be widely conserved. Low minor allele frequencies of large effect SNPs associated with higher tolerance or accumulation of Cd and Hg are consistent with purifying selection against deleterious mutations. Large effect alleles associated with low tolerance or accumulation were more common in our dataset, suggesting excluder alleles may be advantageous. We suspect similar evolutionary principles and processes would apply to other species as well. Our findings therefore suggest functionally conserved roles in these gene families, and potential conservation at macroevolutionary scales in plants, which is important for breeding and transgenic modifications in crop species.

## Author Contributions

TP and JJP conceived of the study. JJP supervised laboratory and phenotypic experiments. MSD and MML conducted phenotypic measurements and data analysis. TP and BH conducted bioinformatics, GWAS, population genomics, and statistical analyses. MRC conducted bioinformatics and statistical analyses. TP and BH drafted the manuscript, with editorial contributions from all authors.

## Acknowledgments

We thank Nevin Young and Roxanne Denny at the University of Minnesota for germplasm, and their support and discussions regarding the Medicago HapMap collection. This work was supported by the Biological and Environmental Research (BER) program, United States Department of Energy. This work was also supported by grants from CSIC (i-LINK1093), MINECO (AGL2013-40758-R) and Agencia Estatal de Investigación, AEI, Spain (AGL2017-88381-R). We thank Kentaro Shimizu for useful discussions and the University of Zurich Functional Genomics Center for computational support. We also thank Gwyneth Halstead-Nussloch for helpful discussion and help with software and scripts.

