## SupplementaryMaterial for "Genome-wide association study reveals complex genetic architecture of cadmium and mercury accumulation and tolerance traits in *Medicago truncatula*"

Supplementary Material

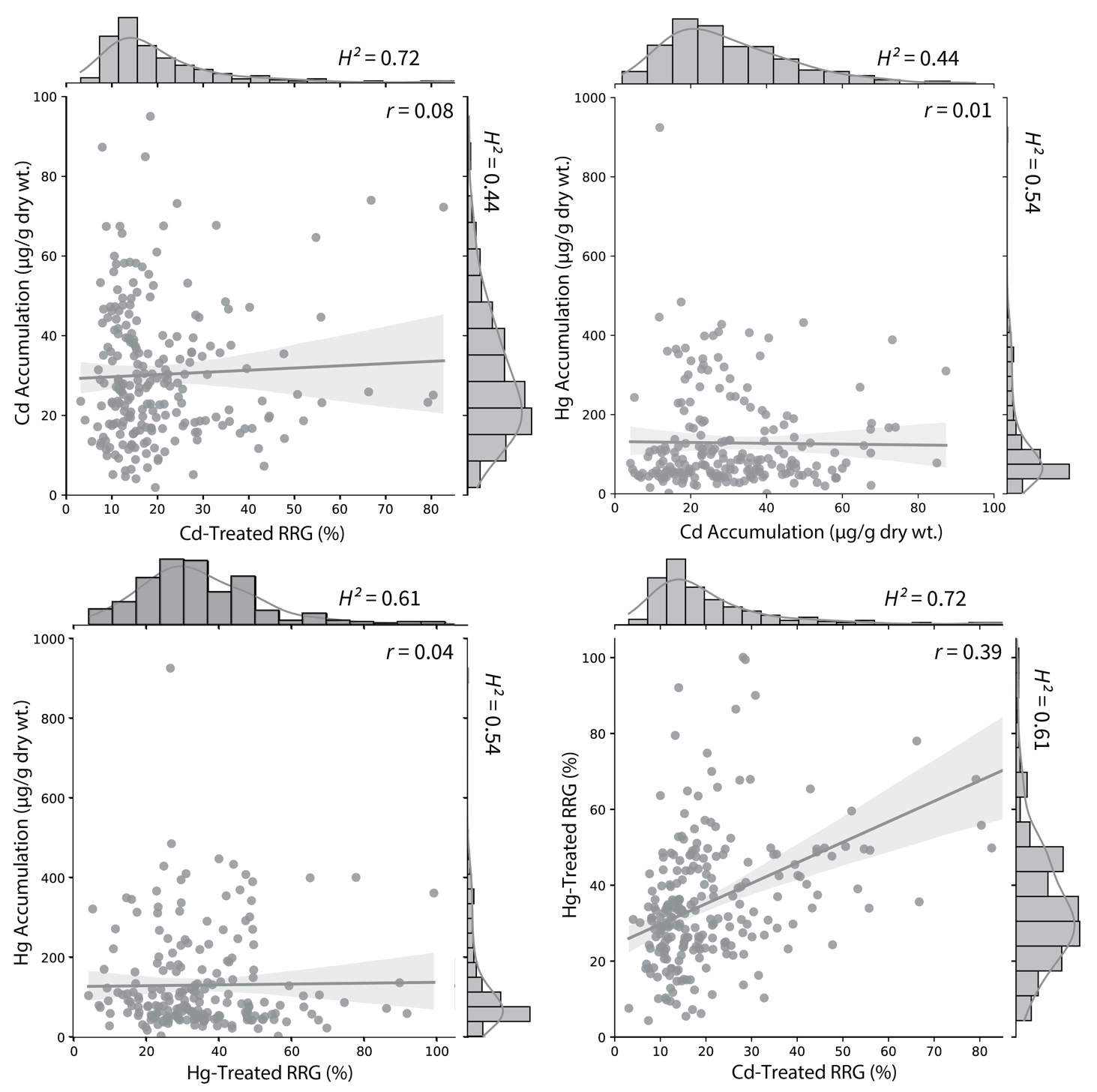

**Supplementary Figure 1.** Phenotypic distributions of leaf accumulation of Cd and Hg in leaves, and relative root growth (RRG) in response to both metals. Corresponding histograms for each phenotype are displayed opposite to the x- and y-axes. Linear regression and 95% confidence intervals are overlayed on the scatter plot. Heritability (*H*^2^) is shown next to each histogram, and Pearson correlation coefficients (*r*) are shown in each panel.

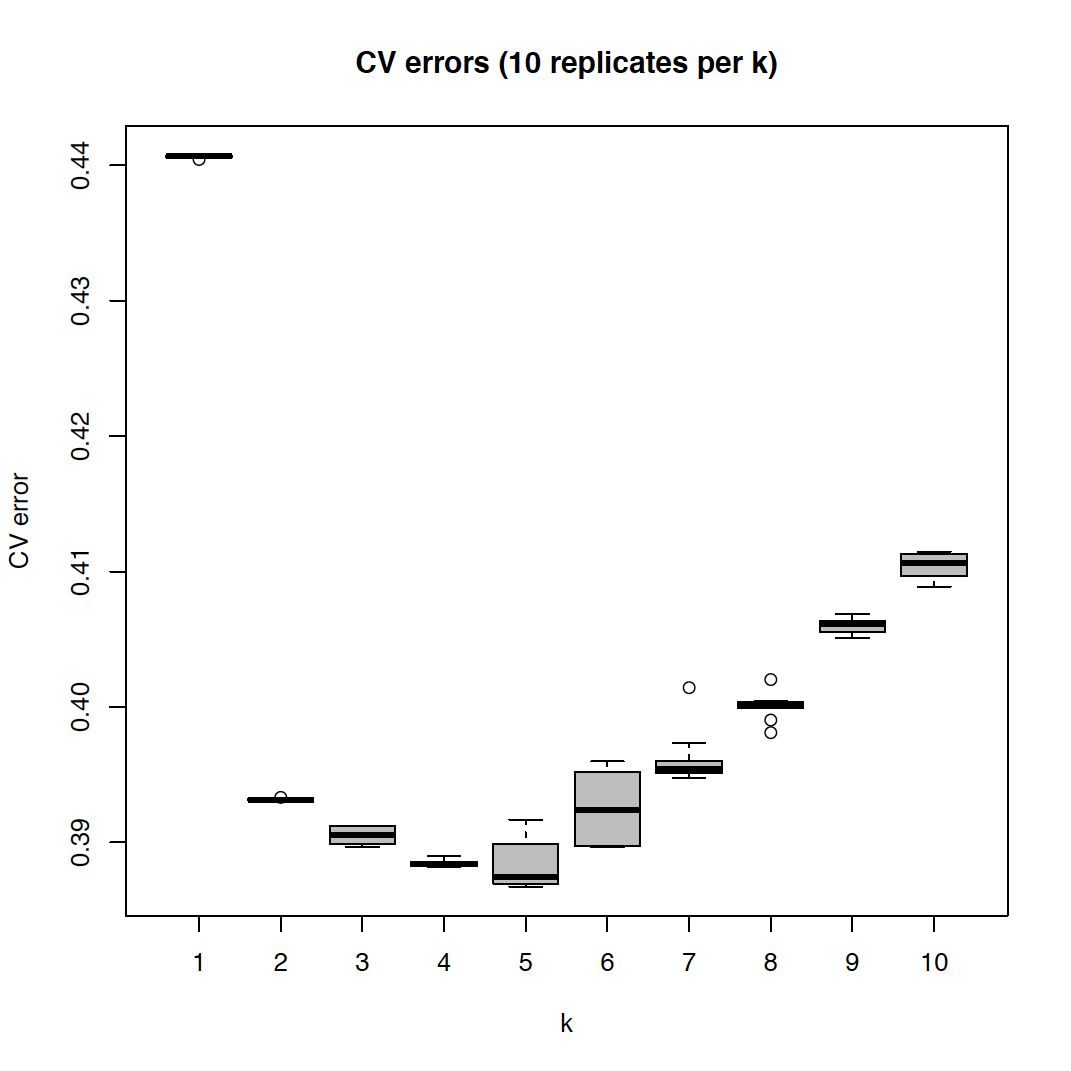

Supplementary Figure 2. Average cross validation errors of 10 Admixture runs for each k between 1 and 10. The y-axis represents the cross-validation error value, the x-axis is k, number of population clusters. The distribution of ancestry components in all samples of the HapMap dataset for the best iteration (lowest cross validation error) was k = 5.

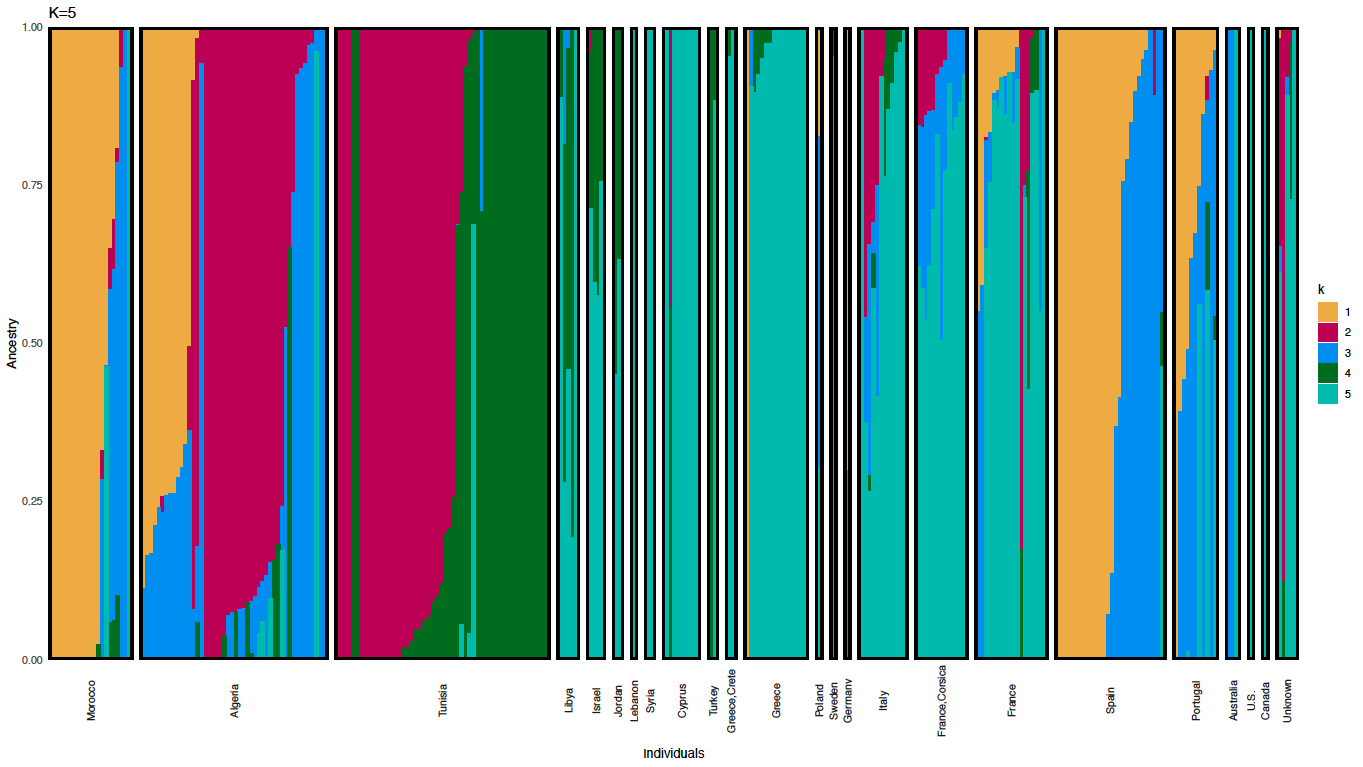

### Supplementary Figure 3. Population structure analysis identified five admixture components. The population structure was determined using the software Admixture. Cross validation errors were lowest for k=5. As expected, genotypes cluster according to geographic location. The most apparent clustering is the East to West separation into k1, k2 and k4. Specifically, k1 comprises of accessions from Spain, Morocco and Algeria, therefore being the west-most cluster. The k2 cluster is oriented toward the East, containing accessions from Algeria and Tunisia, followed by k4, which spans Northeast Africa and Southwest Asia. To a lesser degree a North to South separation is also present, with k3 and k5 containing accessions from regions more to the North than those in k1, k2 or k4. This is especially true for k5, which mostly contains accessions from Greece and Cyprus, therefore being to the North of k4. The distribution of k3 is less clear, but it seems to be represented mostly in Algerian and European accessions and therefore also more to the north. Our structure analysis is comparable to the recent WhoGem analysis by Gentzbittel et al., (2019). Colors represent the 5 components, the y-axis the proportion of each component in an individual and each bar on the x-axis represents an individual.

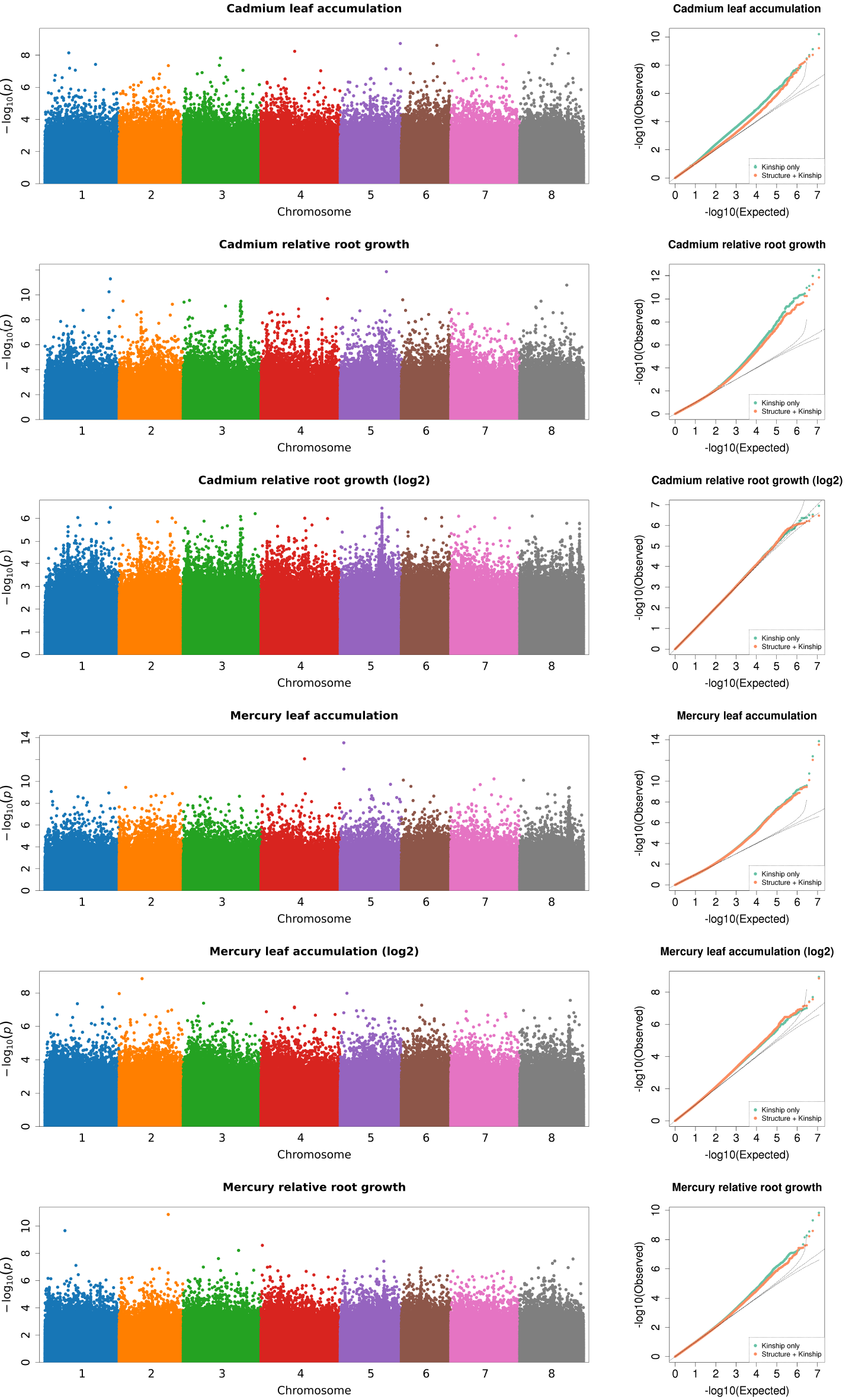

**Supplementary Figure 4**. Genome wide association analysis of heavy metal accumulation and tolerance traits (Manhattan plots shown for each trait; output from GEMMA). Each point represents a SNP at each position in the genome. The number on the x-axis is the chromosome (each color corresponds to one of the eight Mt4.0 chromosomes). The y-axis is the negative log_10_ of the p-values. The higher y-axis values indicate smaller p-values. Next to each Manhattan plot are Quantile-quantile (Q-Q) plots which show the model ﬁt with population structure covariance included (orange points), and without population structure covariance (green points). The x-axis of the Q-Q plots is the expected distribution, the y-axis is the empirical distribution.

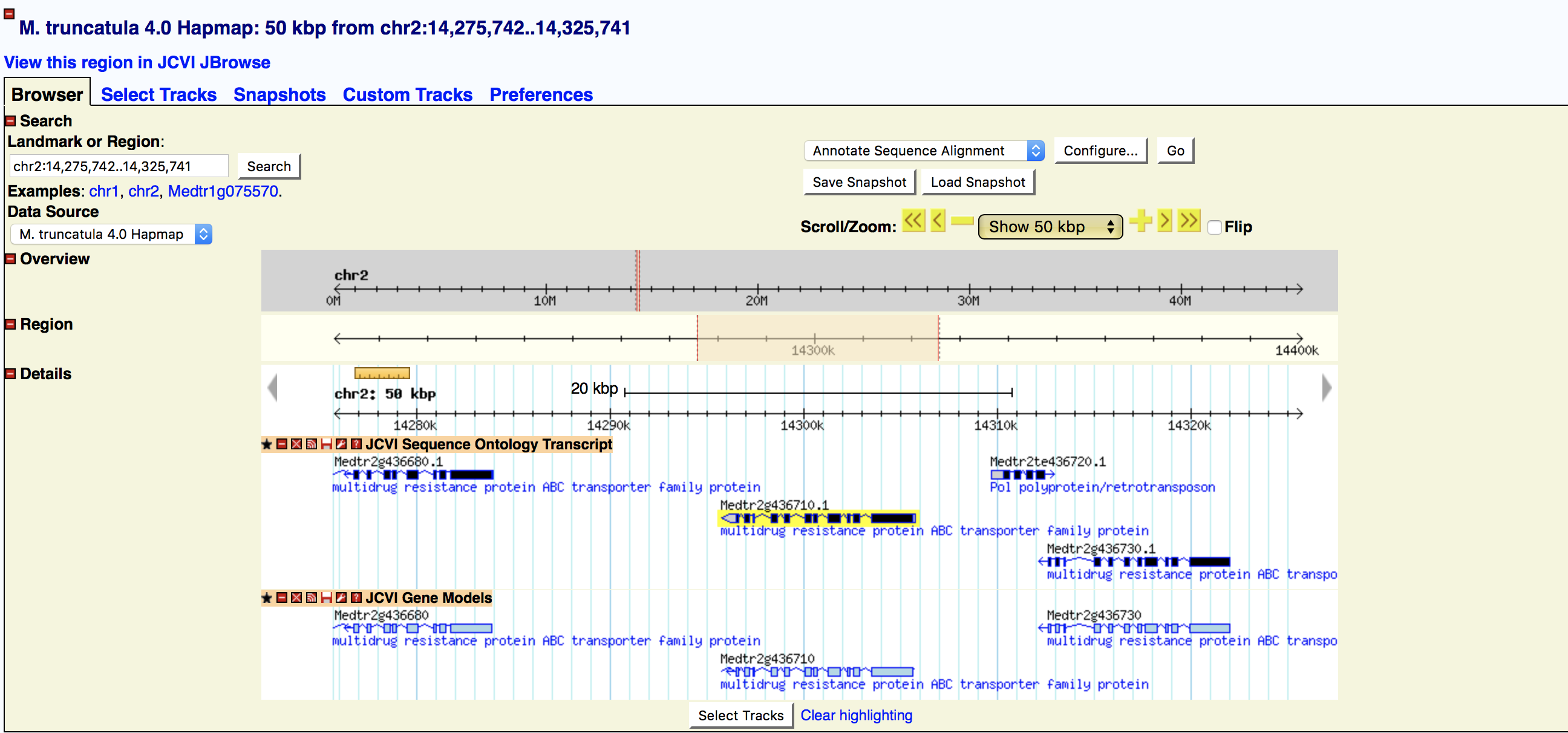

**Supplementary Figure 5.** Genomic organization of three tandem ABC transporters (Medtr2g436680.1, Medtr2g436710, Medtr2g436730) on chromosome 2 in the Mt4.0 reference genome (www.medicagohapmap.org). The closest Blast hit to these three genes is the *Arabidopsis thaliana* ABC transporter *ABCC14* (AT3G62700.1).

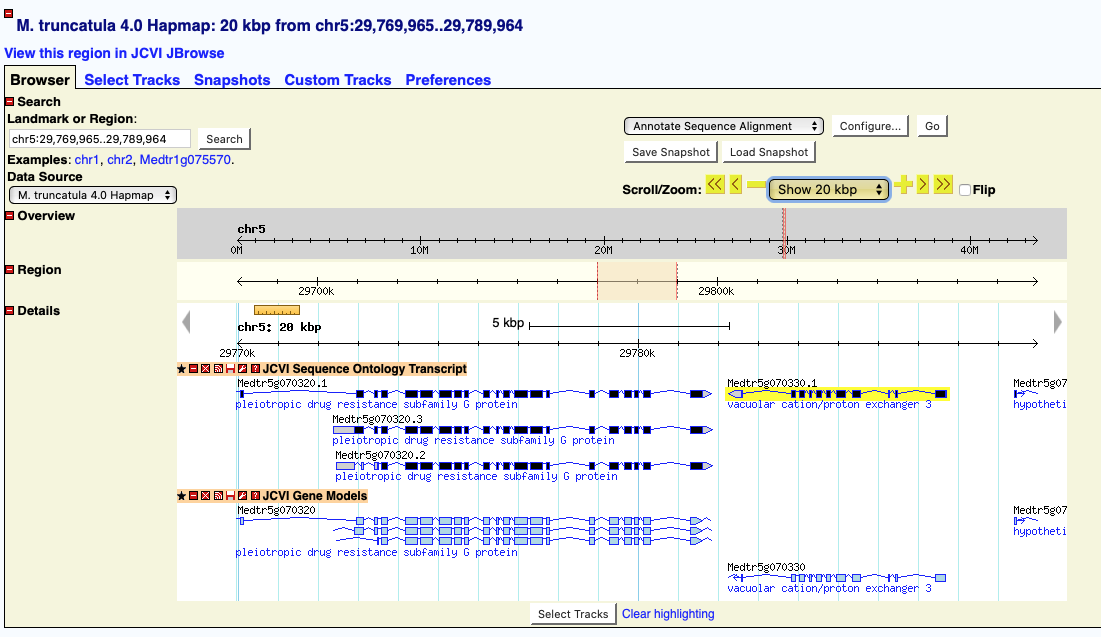

**Supplementary Figure 6**. The Pleiotropic Drug Resistance 3 gene *PDR3* (Medtr5g070320) and Cation Exchanger 3 gene *CAX3* (Medtr5g070330) are 5878 bp apart in the Mt4.0 reference genome (www.medicagohapmap.org).

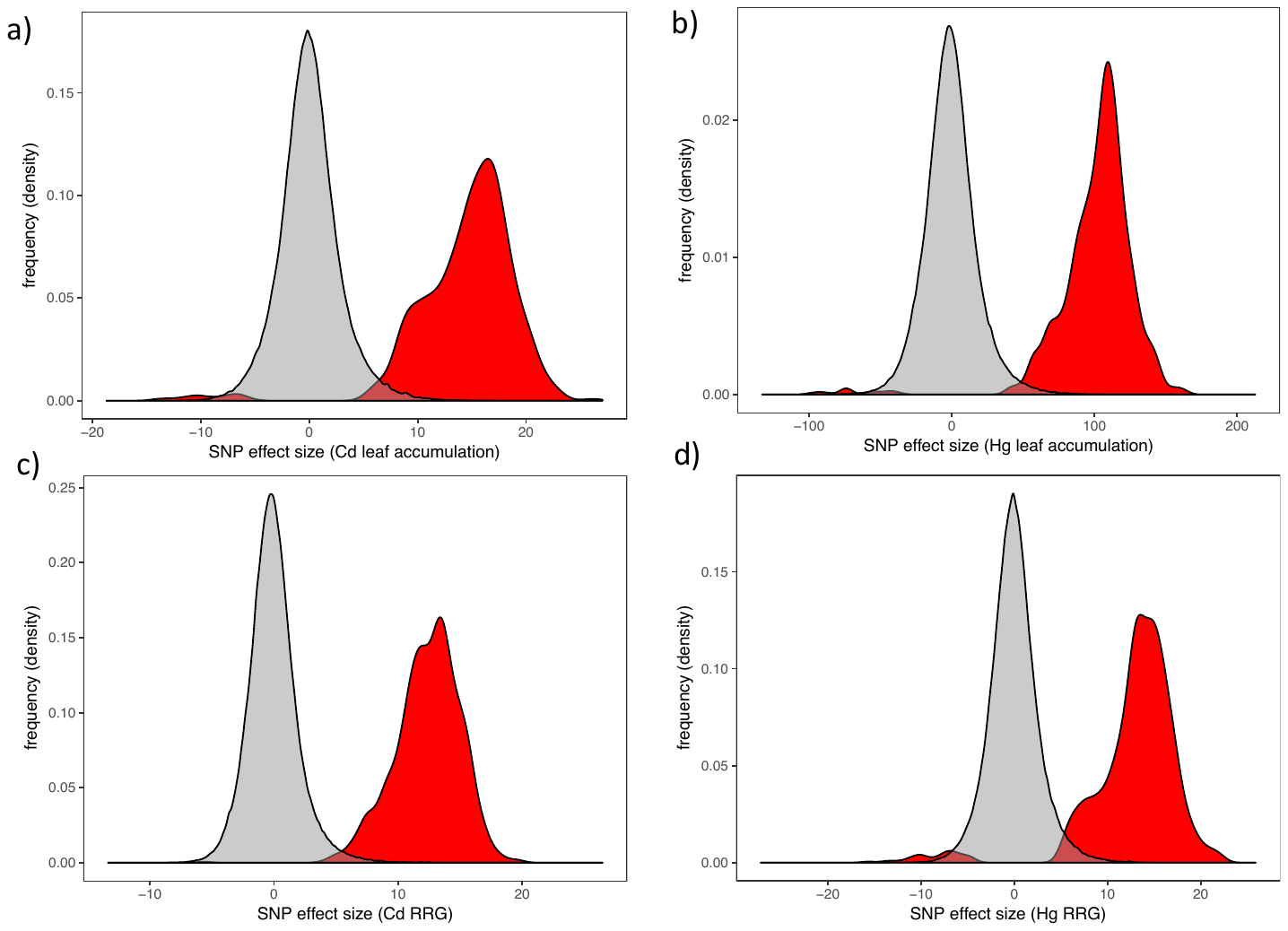

**Supplementary Figure 7.** Density plots showing the genome wide distribution of SNP effect sizes (gray) and the top 1000 SNPs with lowest p-values for each trait (red). The most significant SNPs have the largest effect.

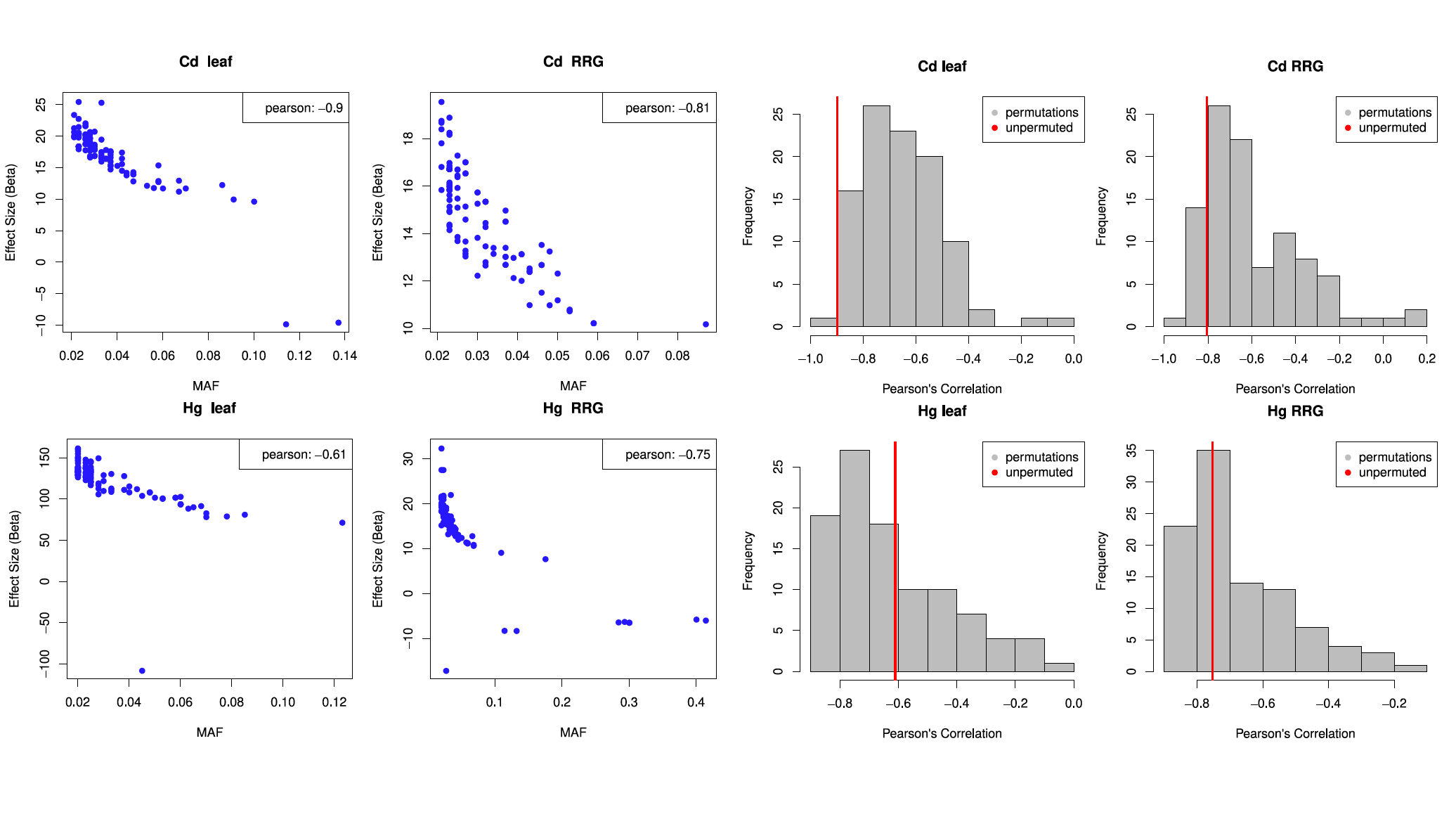

**Supplementary Figure 8.** Effect size and minor allele frequency (MAF) from the top 100 most significant SNPs using the GEMMA package for GWAS (left four panels, one for each trait). Pearson’s correlation coefficients are shown in the upper right corner in each of the four panels. Histograms show the distribution of Pearson’s correlation coefficients from 100 permutations of the GWAS.
